# A *pgr5* suppressor screen uncovers a distinct mechanism safeguarding the cytochrome *b*_6_*f* complex from damage through PGR5

**DOI:** 10.1101/2023.11.28.569138

**Authors:** Jan-Ferdinand Penzler, Belen Naranjo, Sabrina Walz, Giada Marino, Tatjana Kleine, Dario Leister

**Author notes:** These authors contributed equally to this work (Jan-Ferdinand Penzler, Belen Naranjo). Authors for communication. Present address: Instituto de Bioquímica Vegetal y Fotosíntesis, Universidad de Sevilla, Avda. Américo Vespucio 49, 41092 Sevilla, Spain. The author responsible for distribution of materials integral to the findings presented in this article in accordance with the policy described in the Instructions for Authors (https://academic.oup.com/plcell/pages/General-Instructions) is: Dario Leister.

## Abstract

PROTON GRADIENT REGULATION5 (PGR5) is thought to promote cyclic electron flow (CEF) and its deficiency causes increased photosensitivity of photosystem I (PSI), leading to lethality under fluctuating light (FL). By screening for suppressor mutations that rescue FL lethality of *pgr5* plants, we identified a portfolio of mutations affecting 12 photosynthesis-related proteins. Six are required for proper PSII function, one (CcdA) promotes cytochrome (cyt) *b*_6_*f* assembly, and another (PAA1) provides plastocyanin with its copper cofactor. Two other mutations are associated with the chloroplast FBPase cFBP1. This, together with targeted knockout of other genes in the *pgr5* background, suggests three pathways to restore FL viability: (i) reduced electron flow to PSI due to defects in PSII, cyt *b*_6_*f* or plastocyanin but not PSI, (ii) increased electron flow from PSI due to inactivation of ACHT2, a regulator of cFBP1 activity, and (iii) hyperactivity of the NDH-dependent CEF due to inactivation of cFBP1. The remaining two suppressor mutations affected the cyt *b*_6_*f* complex. PFSC1 controls cyt *b*_6_*f* accumulation at early developmental stages. DEIP1/NTA1, previously suggested to be essential for cyt *b*_6_*f* assembly, appears to protect cyt *b*_6_*f* from deleterious effects of PGR5, since plants lacking both DEIP1/NTA1 and PGR5 are viable and accumulate cyt *b*_6_*f*.

## INTRODUCTION

Under natural conditions, plants have to cope with continuous changes in light intensity over the course of the day, for example due to cloud cover and canopy shading. Thus, under low light conditions, maximum light absorption and energy utilization are required to meet metabolic needs through photosynthesis. However, excess light can lead to the production of harmful reactive oxygen species (ROS) and cause irreversible damage to the photosynthetic apparatus or photoinhibition. Therefore, acclimation to light fluctuations is a crucial and adaptive response for photosynthetic organisms to maximize their efficiency in capturing light energy while minimizing the potential damage caused by excessive light exposure (reviewed by: Long et al., 2022; Leister, 2023).

During photosynthesis, electrons are mainly transported from water to the final acceptor NADP^+^ in the thylakoid membranes of chloroplasts via photosystem II (PSII), cytochrome *b*_6_*f* (cyt *b*_6_*f*) and photosystem I (PSI). In the process, a trans-thylakoid proton gradient is generated. The end products are ATP and NADPH, which are required for subsequent reactions. This electron transport is known as linear electron flow (LEF). In addition, other electron transport pathways exist, such as cyclic electron flow (CEF), in which electrons from PSI are transferred back to the plastoquinone (PQ) pool and cyt *b*_6_*f*, maintaining the formation of the proton gradient and thus the synthesis of ATP, but without net production of NADPH (reviewed in: Shikanai and Yamamoto, 2017; Leister, 2020). Hence, the balance between the rates of LEF and CEF is crucial for plant acclimation by adjusting the ATP/NADPH ratio. Furthermore, the proton gradient formed across the thylakoid membrane during both LEF and CEF is required for the activation of a series of protective mechanisms, such as ‘non-photochemical quenching’ (NPQ), which allows the thermal dissipation of the excess absorbed light energy, and the down-regulation of the cyt *b*_6_*f* complex or ‘photosynthetic control’ (reviewed in: Shikanai and Yamamoto, 2017). Consequently, the luminal pH acts as a sensor of excess light, leading to a balance between energy utilization and energy dissipation.

Two types of CEF acting as a Fd-PQ reductases have been described: the NDH-dependent pathway via the NADH dehydrogenase-like complex (NDH), and the PGR5-dependent pathway (also known as the antimycin (AA)-sensitive pathway), which is the dominant and more important pathway in C3 plants (Munekage et al., 2004; Ogawa et al., 2023; reviewed in: Yamori and Shikanai, 2016). ‘Proton Gradient Regulation 5’ (PGR5) is a peripheral thylakoid protein associated with PSI (Munekage et al., 2002) that interacts with and is regulated by the two integral thylakoid proteins ‘PGR5-Like 1’ (PGRL1) (DalCorso et al., 2008) and ‘PGR5-Like 2’ (PGRL2) (Rühle et al., 2021). However, the exact mechanism by which PGR5-dependent CEF takes place is still unclear. Nevertheless, it is evident that in the absence of PGR5, i.e. in *pgr5* and *pgrl1ab* mutants, PSI is inhibited (Munekage et al., 2002; DalCorso et al., 2008; reviewed in: Yamori and Shikanai, 2016).

Defective PGR5-dependent CEF leads to reduced acidification of the lumen, which decreases ATP production that in turn causes overreduction of the stroma due to the reduced ATP/NADPH ratio. It also prevents the induction of NPQ and the downregulation of the cyt *b*_6_*f* complex, leading to an overreduction of PSI. Indeed, plants lacking PGR5 exhibit a pleiotropic phenotype, where the effect on CEF is not easily disentangled from other processes such as stromal overreduction with high PSI acceptor side limitation, and reduced proton motive force due to high ATPase conductance caused by stromal ATP depletion (Avenson et al., 2005; Munekage et al., 2008). Under fluctuating light (FL), this complex defect due to PGR5 deficiency is lethal (Tikkanen et al., 2010; Suorsa et al., 2012), highlighting the importance of this protein for plant survival, as changes in light intensity can occur very rapidly in field canopies. Accordingly, increasing the electron sink capacity of PSI, i.e. decreasing the PSI acceptor side limitation, by adding methyl viologen (Munekage et al., 2002) or overexpressing flavodiiron proteins from *Physcomitrella patens* in Arabidopsis *pgr5* (Yamamoto and Shikanai, 2019) partially rescues the *pgr5* photosynthetic phenotype. In addition, decreased electron transport rates from PSII to PSI, i.e. increased PSI acceptor side limitation, induced either by the addition of DCMU (Suorsa et al., 2012) or the *pgr1* mutation, which affects the cyt *b*_6_*f* complex, also rescues the *pgr5* photosynthetic phenotype (Yamamoto and Shikanai, 2019). Furthermore, combined knock-out of multiple proteins of the oxygen-evolving complex of PSII rescues not only the photosynthetic defect of the *pgr5* mutation but also viability under FL conditions (Suorsa et al., 2016). A third possibility for restoring CEF in the absence of PGR5 is to increase the activity of NDH-dependent CEF, as seen in double mutants of *pgr5* and loss-of-function alleles of the chloroplast FBPase (cFBP1) (Livingston et al., 2010). Taken together, these examples demonstrate that there are multiple interactions between PGR5-dependent CEF, NDH-dependent CEF and other thylakoid electron transport pathways of the thylakoid, although the precise mechanisms remain largely elusive.

In this context, the cyt *b*_6_*f* complex occupies a key position between the two photosystems (PSII and PSI), taking part in both LEF and CEF. The structure of the dimeric complex has been determined at high resolution from several species (Zhang et al., 2023). However, relatively little is known about its assembly, degradation and interaction with stromal and thylakoid components involved in CEF. Indeed, a new component involved in the assembly of the cyt *b*_6_*f* has recently been identified, DEIP1/NTA1 (Sandoval-Ibanez et al., 2022; Li et al., 2023). The cyt *b*_6_*f* complex is the site of proton translocation across the thylakoid membrane, establishing the proton gradient required for ATP synthesis and photoprotection, and an important component of the aforementioned photosynthetic control. Additionally, cyt *b*_6_*f* is involved in the activation of the so-called “state transitions”, which balance the excitation energy between PSII and PSI through phosphorylation and relocation of the antenna complex LHCII (reviewed in: Malone et al., 2021). Therefore, it is not surprising that cyt *b*_6_*f* has emerged as a regulatory hub for the coordination of different electron transport pathways and mechanisms for short- and long-term light acclimation (Yamamoto and Shikanai, 2019). In fact, the finding that the *pgr1* mutation, conferring hypersensitivity of the cyt *b*_6_*f* complex to luminal acidification, rescues the *pgr5* photosynthetic phenotype (Yamamoto and Shikanai, 2019), raises the question of the extent to which the cyt *b*_6_*f* complex and PGR5-dependent processes functionally interact.

With the aim of identifying new components involved in FL acclimation, as well as deepening our knowledge of the mechanisms and the interplay between the different photosynthetic electron transport pathways, we have performed a *pgr5* suppressor screen based on the lethal phenotype of the Arabidopsis *pgr5* mutant under FL. We selected 449 *pgr5* suppressor mutants, of which we sequenced 37 and identified 25 mutations affecting 12 different photosynthesis-related proteins, including a novel one, PFSC1, which controls cyt *b*_6_*f* accumulation at early developmental stages, and two mutations affecting the functionality or regulation of cFBP1. We demonstrated that changes in PSI donor and acceptor side limitations at early developmental stages are responsible for the recovery of viability of young *pgr5* plants under FL light. Moreover, we found a completely unexpected interplay between PGR5 and DEIP1/NTA1, revising the concept that DEIP1/NTA1 acts as a cyt *b*_6_*f* assembly factor and suggesting a link between PGR5 function and cyt *b*_6_*f* accumulation.

## RESULTS

### A *pgr5* suppressor screen based on recovery of viability under fluctuating light

To identify novel components involved in light acclimation, we performed a *pgr5* suppressor screen under FL conditions. After mutagenizing *pgr5* Arabidopsis seeds using EMS, we screened 273.000 plants in the second generation (M_2_) to saturate the screen and selected 449 survivors under FL in contrast to *pgr5*, which was lethal after the cotyledon stage in this condition. When we obtained the genome sequences of the first four suppressor lines, designated as *p5s1*, *2*, *3,* and *4*, we found mutations affecting PSII function in all of them, in agreement with the results of Suorsa et al. (2016). Therefore, in order to eliminate most mutations in PSII components, we performed an additional selection step and considered for further analysis only those lines with maximum PSII quantum yield (F*v*/F*m*) values similar to wild type (WT) (117 lines). Finally, 37 *pgr5* suppressor mutants were sequenced and analyzed for causative mutations. In 25 cases, we identified single nucleotide polymorphisms (SNPs) affecting 12 different genes as potential causative mutations of the *pgr5* suppressor phenotype (**Table 1**). Five genes were hit by independent mutations, showing that the screen was saturated and providing evidence that those mutations were indeed responsible for the suppressor phenotype. We confirmed 6 mutations, in particular those for which we obtained only one mutant allele during the screen, by identifying or generating additional mutant alleles (T-DNA knockouts or CRISPR-Cas induced mutations) and introducing them into the *pgr5* background to verify the non-lethal phenotype under FL (*pgr5 lpa66* for *p5s1* and *p5s2*, *pgr5 mterf5* for *p5s3*, *pgr5 cfbp1* for *p5s18* and *p5s19* and *pgr5 at2g04360* for *p5s22*, *p5s23*, *p5s24* and *p5s25*) or CRISPR-Cas9 (*pgr5 acht2* for *p5s20* and *pgr5 deip1* for *p5s21*) (**Table 1**, see **Figure 1A** for FL phenotype of representative lines).

**Figure 1.**
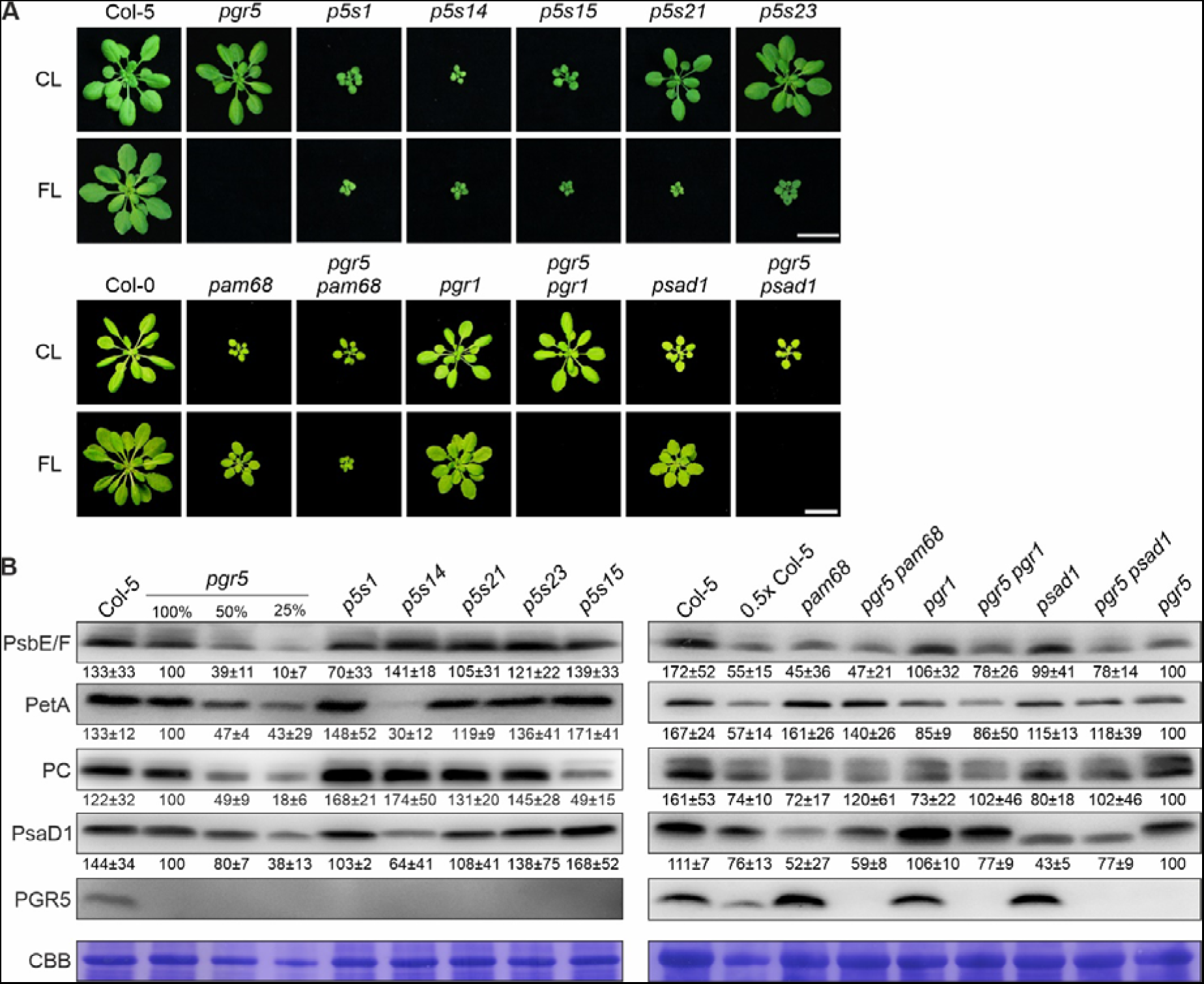
Impact of perturbed linear electron flow on the suppression of *pgr5* phenotypes. **(A)** Growth phenotype of suppressor lines (*p5s1*, *p5s14*, *p5s15*, *p5s21* and *p5s23*) and representative affecting PSII (*pam68*), cyt *b*_6_*f* (*pgr1*) and PSI (*psad1*) in either WT or *pgr5* background, compared to *pgr5* and WT (Col-0 and Col-5) under control (CL, after 3 weeks) and fluctuating light (FL, after 5 weeks). Scale bars indicate 1 cm. **(B)** Immunoblot analysis of representative photosynthetic proteins. Aliquots of total leaf protein from the CL-grown plants shown in (A) were fractionated by SDS-PAGE under reducing conditions, transferred onto PVDF membranes, and immunodecorated with specific antibodies. Staining of the membranes with Coomassie Brilliant Blue (C.B.B.) served as loading control. Representative blots from three experiments are presented, as well as the average of the band quantifications (n=3) relative to *pgr5* (100%) ± SD.

**Table 1.**
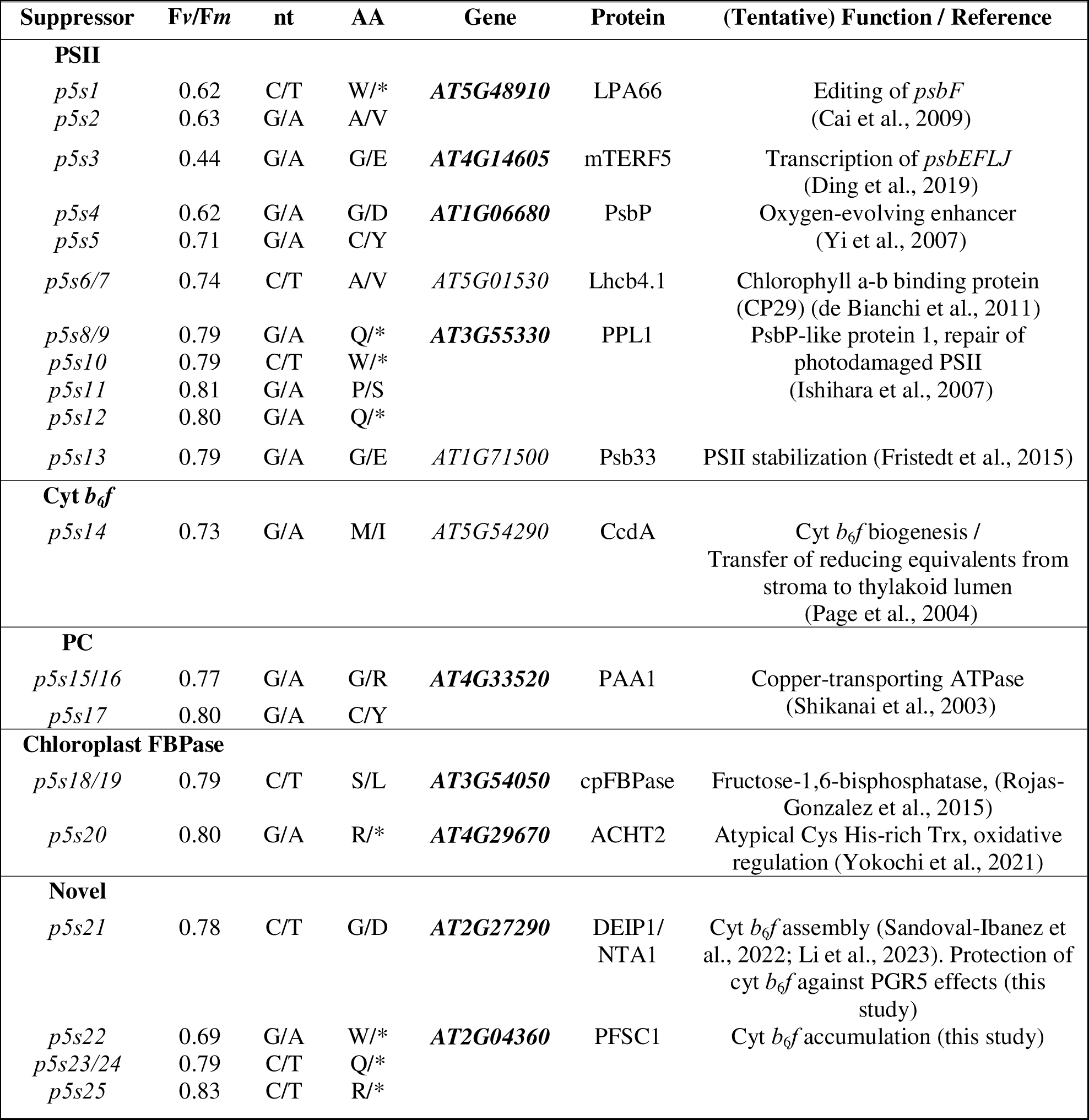
Overview of the *pgr5* suppressors identified in the screen. . The maximum quantum yield of PSII (F*v*/F*m*), single nucleotide polymorphisms (SNPs, nt), corresponding amino acid exchanges (AA), ATG accession number, protein name and its tentative function together with relevant literature are listed. Genes for which more than one independent suppressor mutation was found or whose suppressor phenotype was confirmed by T-DNA or CRISPR/Cas mutant alleles are highlighted in bold.

| Suppressor | Fv/Fm | nt | AA | Gene | Protein | (Tentative) Function / Reference |
| --- | --- | --- | --- | --- | --- | --- |
| **PSII** |  |  |  |  |  |  |
| *p5s1* | 0.62 | C/T | W/* | **AT5G48910** | LPA66 | Editing of *psbF* (Cai et al., 2009) |
| *p5s2* | 0.63 | G/A | A/V |  |  |  |
| *p5s3* | 0.44 | G/A | G/E | **AT4G14605** | mTERF5 | Transcription of *psbEFLJ* (Ding et al., 2019) |
| *p5s4* | 0.62 | G/A | G/D | **AT1G06680** | PsbP | Oxygen-evolving enhancer (Yi et al., 2007) |
| *p5s5* | 0.71 | G/A | C/Y |  |  |  |
| *p5s6/7* | 0.74 | C/T | A/V | *AT5G01530* | Lhcb4.1 | Chlorophyll a-b binding protein (CP29) (de Bianchi et al., 2011) |
| *p5s8/9* | 0.79 | G/A | Q/* | **AT3G55330** | PPL1 | PsbP-like protein 1, repair of photodamaged PSII (Ishihara et al., 2007) |
| *p5s10* | 0.79 | C/T | W/* |  |  |  |
| *p5s11* | 0.81 | G/A | P/S |  |  |  |
| *p5s12* | 0.80 | G/A | Q/* |  |  |  |
| *p5s13* | 0.79 | G/A | G/E | *AT1G71500* | Psb33 | PSII stabilization (Fristedt et al., 2015) |
| **Cyt *b<sub>6</sub>f*** |  |  |  |  |  |  |
| *p5s14* | 0.73 | G/A | M/I | *AT5G54290* | CcdA | Cyt *b<sub>6</sub>f* biogenesis / Transfer of reducing equivalents from stroma to thylakoid lumen (Page et al., 2004) |
| **PC** |  |  |  |  |  |  |
| *p5s15/16* | 0.77 | G/A | G/R | **AT4G33520** | PAA1 | Copper-transporting ATPase (Shikanai et al., 2003) |
| *p5s17* | 0.80 | G/A | C/Y |  |  |  |
| **Chloroplast FBPase** |  |  |  |  |  |  |
| *p5s18/19* | 0.79 | C/T | S/L | **AT3G54050** | cpFBPase | Fructose-1,6-bisphosphatase, (Rojas-Gonzalez et al., 2015) |
| *p5s20* | 0.80 | G/A | R/* | **AT4G29670** | ACHT2 | Atypical Cys His-rich Trx, oxidative regulation (Yokochi et al., 2021) |
| **Novel** |  |  |  |  |  |  |
| *p5s21* | 0.78 | C/T | G/D | **AT2G27290** | DEIP1/NTA1 | Cyt *b<sub>6</sub>f* assembly (Sandoval-Ibanez et al., 2022; Li et al., 2023). Protection of cyt *b<sub>6</sub>f* against PGR5 effects (this study) |
| *p5s22* | 0.69 | G/A | W/* | **AT2G04360** | PFSC1 | Cyt *b<sub>6</sub>f* accumulation (this study) |
| *p5s23/24* | 0.79 | C/T | Q/* |  |  |  |
| *p5s25* | 0.83 | C/T | R/* |  |  |  |

### Perturbation of linear electron flow at PSII and the cyt *b*_6_*f* complex, but not PSI, can rescue *pgr5* lethality

Interestingly, all the SNPs identified in the screen affected genes encoding proteins involved in photosynthesis (**Table 1**). Six genes are required for proper PSII function, such as *LPA66* (mutated in *p5s1* and *p5s2*), which is required for PSII assembly by editing *psbF* (Cai et al., 2009). Another gene, *CcdA*, mutated in *p5s14*, is necessary for the biogenesis of cyt *b*_6_*f* (Page et al., 2004). The two suppressor lines *p5s15* and *p5s17* carry mutations in the gene for the copper transporter PAA1, which is required for the accumulation of functional plastocyanin (PC) (Shikanai et al., 2003). Under control light (CL; 12 h light with 100 µmol photons m^−2^ s^−1^ / 12 h darkness, see Methods) conditions, *p5s1*, *14* and *15* showed markedly reduced growth compared to *pgr5* and WT, and they survived under FL in contrast to *pgr5* (**Figure 1A**).

In order to clarify the effects of these representative mutations on the levels of the corresponding photosynthetic multiprotein complexes (*p5s1* and *p5s14*) and PC (*p5s15*), immunoblot analyses were performed. Indeed, compared to *pgr5* plants, *p5s1, p5s14* and *p5s15* showed only 70% of PsbE/F, 30% of PetA, and 49% of PC levels, respectively (**Figure 1B**). This supports the previous finding of Suorsa et al. (2016) that defects in PSII can suppress *pgr5* lethality under FL conditions. It adds defects in the cyt *b*_6_*f* complex and plastocyanin accumulation to this repertoire of *pgr5* lethality suppressors, prompting the idea that an increase in PSI donor side limitation may be the common mechanism behind this.

In addition, we identified a SNP in a gene encoding a protein recently described to be involved in the assembly of the cyt *b*_6_*f* complex and essential for photoautotrophic growth, DEIP1/NTA1 (Sandoval-Ibanez et al., 2022; Li et al., 2023). Interestingly, the corresponding suppressor line *p5s21* carrying a mutation in DEIP1/NTA1 was able to grow similarly to the WT under CL conditions (**Figure 1A**) and had WT-like PetA levels (**Figure 1B**). This finding is surprising and deserves special attention because the *deipt1*/*nta1* mutation alone causes lethality at the seedling stage (Sandoval-Ibanez et al., 2022; Li et al., 2023).

Furthermore, three independent suppressor lines were found (*p5s22*, *p5s23*/*24* and *p5s25*) which carry mutations in the *AT2G04360* gene encoding an unknown protein. Analysis of *p5s23* showed that under CL conditions the mutant grew like the WT and had similar or even higher levels of all thylakoid proteins tested (**Figure 1A,B**). We provide a detailed characterization of the *p5s21* and *p5s23* lines and their effects on the DEIP1/NTA1 and At2g04360 proteins later in the manuscript.

Our suppressor screen failed to identify mutations in PSI-related genes or in *PETC*/*PGR1*, suggesting that our screen was either not saturated or that defects in PSI and PGR1 cannot suppress *pgr5* lethality. To clarify this, we generated a series of double mutants in the *pgr5* background. As expected from the results of the screen and previous findings (Suorsa et al., 2016), mutation of the PSII assembly factor PAM68 (Armbruster et al., 2010) suppressed the *pgr5* phenotype under FL (**Figure 1A**). However, mutation of *PSAD1*, which affects accumulation of the D-subunit of PSI (Ihnatowicz et al., 2004), failed to suppress *pgr5,* consistent with our negative results for PSI-related genes from the suppressor screen. Interestingly, *pgr5 pgr1*, which lacks PGR5 and contains a mutant Rieske protein in cyt *b*_6_*f* that does not alter protein composition but reduces cyt *b*_6_*f* activity under high light (Munekage et al., 2001)(**Figure 1B**), did not suppress *pgr5* lethality under FL (**Figure 1A**). This indicates that the previously observed increase in photosynthetic control associated with increased PSI donor side limitation due to the *pgr1* mutation (Yamamoto and Shikanai, 2019) is not sufficient to rescue photoautotrophic growth of young plants under FL conditions.

### Mutual compensation of defects caused by *pgr5* and mutations affecting cFBP1 activity

The two suppressor lines *p5s18/19* and *p5s20* carry mutations in the *CFBP1* gene, which encodes cFBP1, and the *ACHT2* gene. The ACHT2 protein is involved in cFBP1 oxidation, simultaneous inactivation of ACHT1 and ACHT2 can suppress *ntrc* phenotypes, and ACHT2 overexpression results in higher NPQ, stunted growth, lower chlorophyll and F*v*/F*m* (Yokochi et al., 2021). Under CL conditions, the two suppressor lines grew like WT, and they survived under FL in contrast to *pgr5* (**Figure 2A**). To confirm that the suppression of the *pgr5* mutation was indeed caused by the two mutations, we identified loss-of-function alleles of *CFPB1* and *ACHT2* and introduced them into the *pgr5* background. We generated a double mutant line (*pgr5 cfbp1-1*) by crossing *pgr5* with a T-DNA insertion line lacking detectable expression of cFBP1 (*cfbp1-1*) (**Figure 2B**). The growth of *cfbp1-1* under CL conditions was severely impaired, as previously reported (Livingston et al., 2010; Rojas-Gonzalez et al., 2015). However, growth was restored to WT levels in the *pgr5 cfbp1-1* and *p5s18/19* lines, an observation that went unnoticed by Livingston et al. (2010), who also generated such double mutants. Since *cfbp1-1* plants were found to have significantly and constitutively increased NDH-dependent CEF (Livingston et al., 2010), they were also named HCEF1 for “high CEF1”. Our finding that depletion of PGR5-CEF in this background of constitutive and enhanced NDH-CEF restores normal growth sheds new light on the original findings of Livingston et al. (2010), as it provides an indirect support for the paradigm that PGR5 is indeed primarily involved in CEF. Taken together with the finding that increased PGR5-CEF via PGR5 overexpression has detrimental effects on plant growth (Okegawa et al., 2007; Long et al., 2008), the most parsimonious explanation for the suppression of the *cfbp1-1*/*hcef1*-related growth phenotype by *pgr5* is that the depletion of PGR5-CEF balances the overall level of CEF to WT-like levels. In fact, this compensatory response is already found in the *cfbp1-1* line, where PGR5 accumulation is markedly decreased (**Figure 2B**), as previously observed (Livingston et al., 2010).

**Figure 2.**
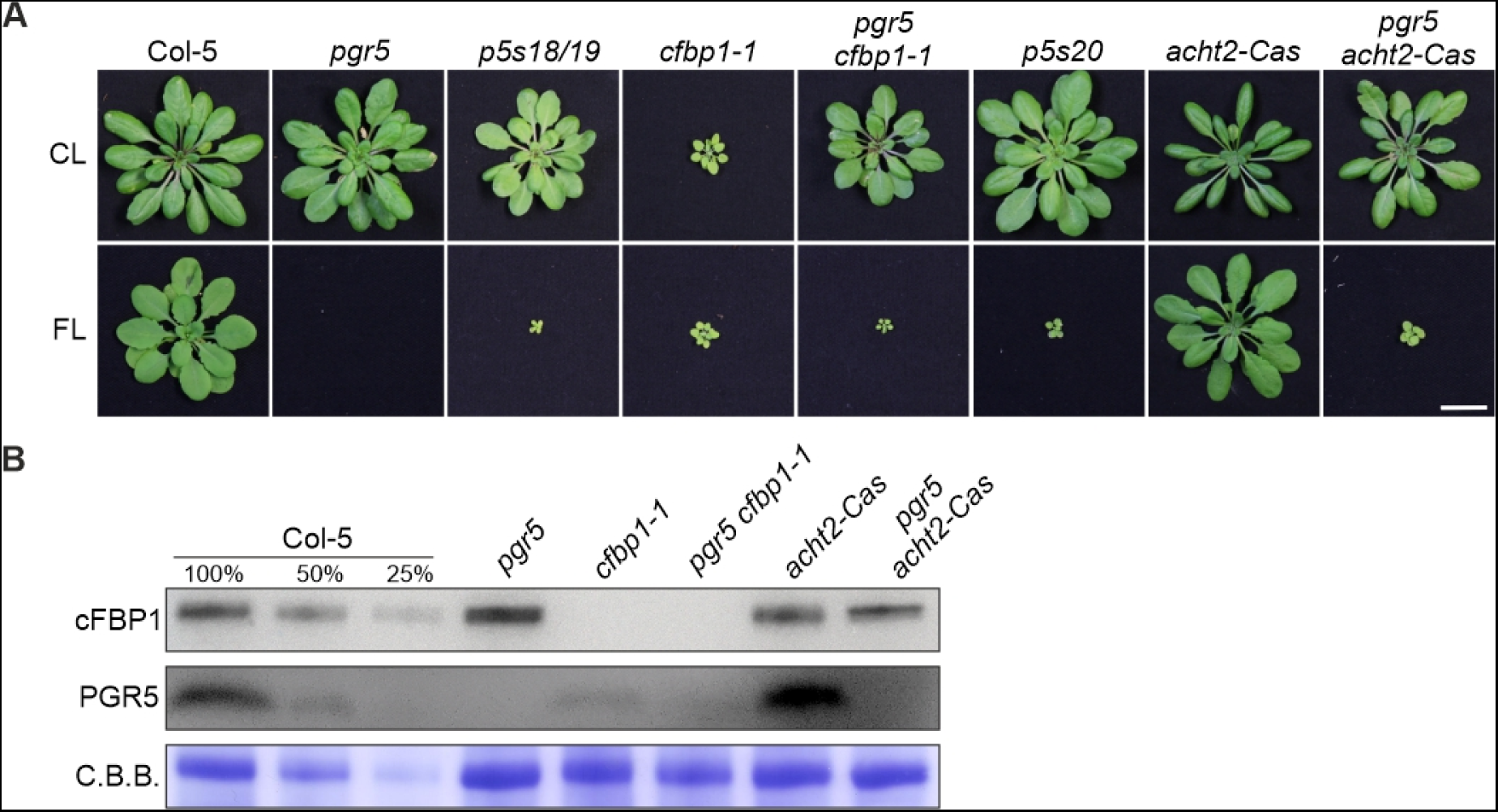
Effect of loss or altered regulation of cFBP1 on the suppression of *pgr5* phenotypes. **(A)** Growth phenotype of WT (Col-5), single (*pgr5*, *cfbp1-1* and *acht2-Cas*) and double mutant (*pgr5 cfbp1-1* and *pgr5 acht2-Cas*) plants under control (5 weeks CL) and FL (5 weeks) conditions. Scale bar indicates 1 cm. **(B)** Aliquots of total leaf proteins were isolated from the CL-grown plants in (A), fractionated by SDS-PAGE under reducing conditions, transferred onto PVDF membranes, and immunodecorated with cFBP1- or PGR5-specific antibodies. Staining of the membranes with Coomassie Brilliant Blue (C.B.B.) served as loading control.

Moreover, using the CRISPR/CAS system, we generated two knock-out lines for *ACHT*2: one in the Col-0 background (*acht2-Cas*), and the other in *pgr5* (*pgr5 acht2-Cas*) (**Supplemental Figure S1A**). Notably, and similar to the case of the cFBP1 mutant described above, *pgr5 acht2-Cas* displayed the same FL phenotype as *p5s20* (**Figure 2A**).

In conclusion, inactivation of cFBP1 or of its negative regulator can restore the viability of *pgr5* plants under FL conditions. In the case of *pgr5 cfbp1-1*, the *pgr5* mutation suppresses the growth defect of *cfbp1* lines, indicating that the total level of CEF, including PGR5- and NDH-CEF, is critical for plant development.

### Relationship between *pgr5* suppression and changes in PSI donor or acceptor side limitation

To analyze photosynthetic performance, we measured chlorophyll fluorescence of adult plants under conditions mimicking the FL of the screen (for a time-resolved example of values obtained for WT and *pgr5*, see **Supplemental Figure S2**). This analysis showed that during the low light (LL) phase, the three double mutants *p5s1*, *14* and *15* had lower photosynthetic electron transport rates from PSII (Y(II)) and PSI (Y(I)) and an increased deficit of electron donors to PSI (Y(ND)), together with an increase in NPQ, compared to *pgr5* plants (**Figure 3A**). In contrast, during the high light (HL) phase, *p5s1*, *14* and *15* showed slightly lower Y(II) but slightly higher Y(I) values, while Y (ND) was massively increased at the expense of a much less limited PSI acceptor side (Y(NA)) compared to *pgr5* (**Figure 3B**). This suggests that the impaired electron low due to impaired PSII (*p5s1*), cyt *b*_6_*f* (*p5s14*) or PC (*p5s15*) is indeed responsible for suppressing the lethality of the *pgr5* mutation. Indeed, the *pam68* mutation in the *pgr5* background mimics the behavior observed in *p5s1*, whereas neither the *psad1* nor the *pgr1* mutation can markedly reduce the PSI acceptor side limitation of *pgr5* during the HL phase, providing an explanation for their inability to suppress the FL lethality of *pgr5*.

**Figure 3.**
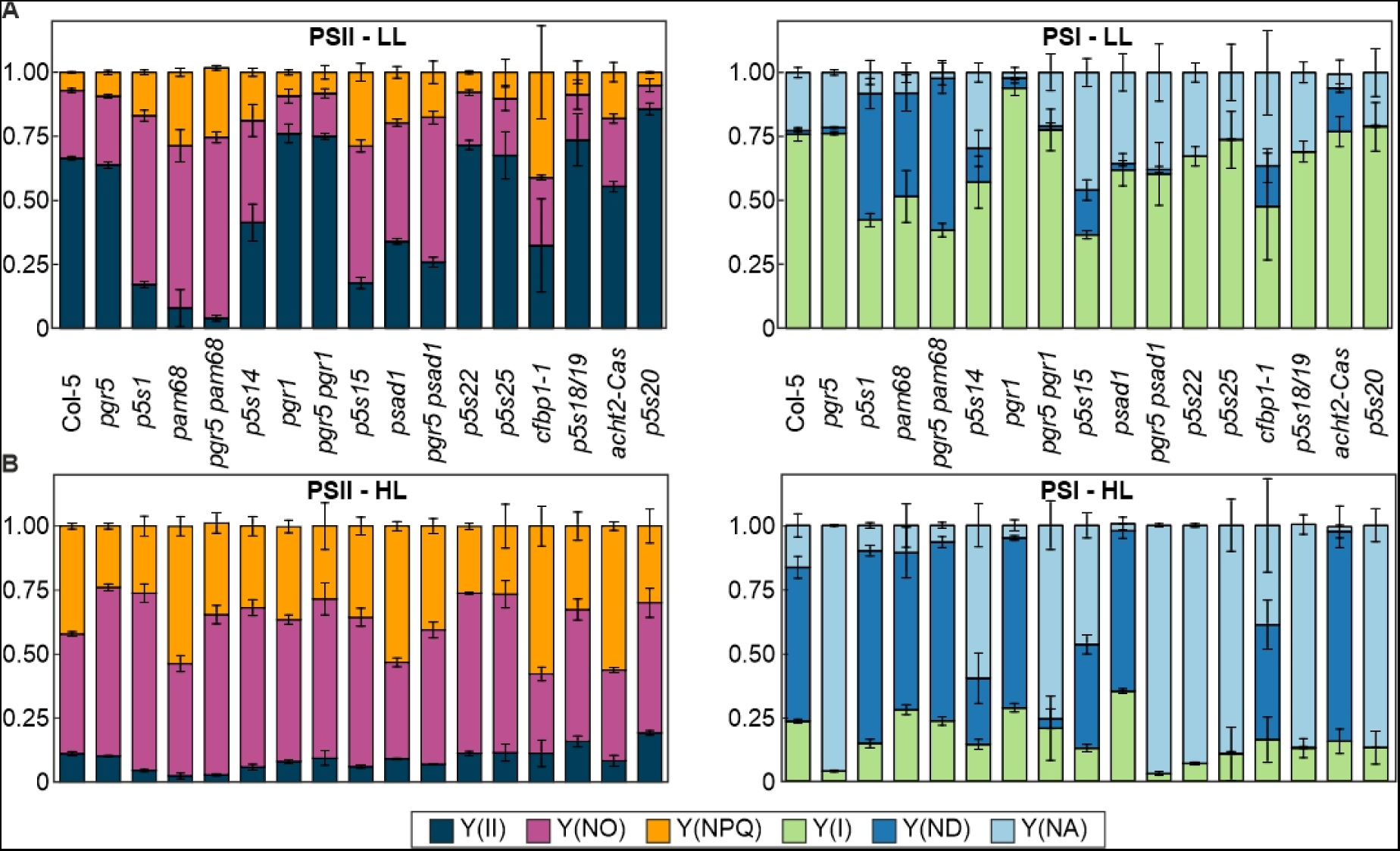
Relationship between *pgr5* suppression and changes in photosynthetic electron flow. **(A,B)** The same genotypes as in Figure 1 were grown under control light for 5 weeks and then subjected to a FL program using the DUAL-PAM chlorophyll fluorometer, applying cycles of 5 min low (50 µmol photons m^−2^ s^−1^) and 1 min high (500 µmol photons m^−2^ s^−1^) actinic light. The values corresponding to PSII (left) and PSI (right) parameters during a low light (LL) (**A**) and high light phase (**B**) are average values of at least 3 replicates ± SD. Y(II), PSII quantum yield; Y(NO), non-regulated energy dissipation, Y(NPQ), non-photochemical quenching; Y(I), PSI quantum yield; Y(ND), PSI donor-side limitation; Y(NA), PSI acceptor-side limitation.

Indeed, the ability of mutations to decrease under HL the PSI acceptor side limitation while increasing the donor side limitation is a feature of the *pgr5* suppressor lines that are affected in LEF between PSII and PSI. However, neither the two suppressor lines with mutations associated with cFBP1 (*p5s18/19* and *p5s20*), nor the two lines with mutations in DEIP1/NTA1 (*p5s21*) or At2g04360 (*p5s23*) differ significantly from *pgr5* with respect to their Y(ND) and Y(NA) values. As the photosynthesis measurements could only be performed on adult plants for technical reasons, since only mature leaves can be measured for this set of parameters, a plausible explanation is that these parameters are indeed altered in younger plants, where they are crucial for survival under FL. In fact, the in-depth analysis of two suppressor lines with mutations in DEIP1/NTA1 (*p5s21*) or At2g04360 (*p5s23*) described below confirms this hypothesis.

### Control of cyt *b*_6_*f* accumulation by DEIP1/NTA1 depends on PGR5 and is developmentally regulated

The *pgr5* suppressor line *p5s21* contained a C/T mutation that resulted in an amino acid exchange (G72D) in the soluble N-terminus of the DEIP1/NTA1 protein (**Figure 4A** and **Supplemental Figure S2B**). To prove that this mutation was responsible for the recovery of *pgr5* viability under FL (**Figure 1A**), we generated two independent *pgr5 deip1/nta1* mutants using CRISPR-Cas technology: *pgr5 deip1-Cas#1* and *#2* (**Figure 4A** and **Supplemental Figure S2B**). It is worth noting that the same mutations could not be obtained in the WT background, which is consistent with previous findings reporting that the single mutant *deip1*/*nta1* is lethal, because the plant does not accumulate cyt *b*_6_*f* (Sandoval-Ibanez et al., 2022; Li et al., 2023). Importantly, the two generated *pgr5 deip1-Cas* lines behaved similar to *p5s21* and were able to grow under FL (**Figure 4B and C**). In both CIRSPR-Cas lines, the DEIP1/NTA1 protein and PGR5 were missing (**Figure 4C**), whereas in *p5s21* residual amounts of DEIP1/NTA1 were detectable. This suggests that the amino acid exchange G72D inactivates DEIP1/NTA1 but does not abolish the accumulation of the inactive protein. Interestingly, in the two *pgr5 deip1-Cas* double mutants, the levels of cyt *b*_6_*f* proteins (PetA, PetB, PetC) were similar to those in *pgr5* and WT plants (**Figure 4C**), in contrast to single *deip1/nta1* mutants that lack cyt *b*_6_*f* (Sandoval-Ibanez et al., 2022; Li et al., 2023). These results suggest that PGR5 plays a relevant role in the mode of action of DEIP1/NTA1 and in the stability of cyt *b*_6_*f*. However, they do not explain the survival of the *pgr5 deip1*/*nta1* plants under FL.

**Figure 4.**
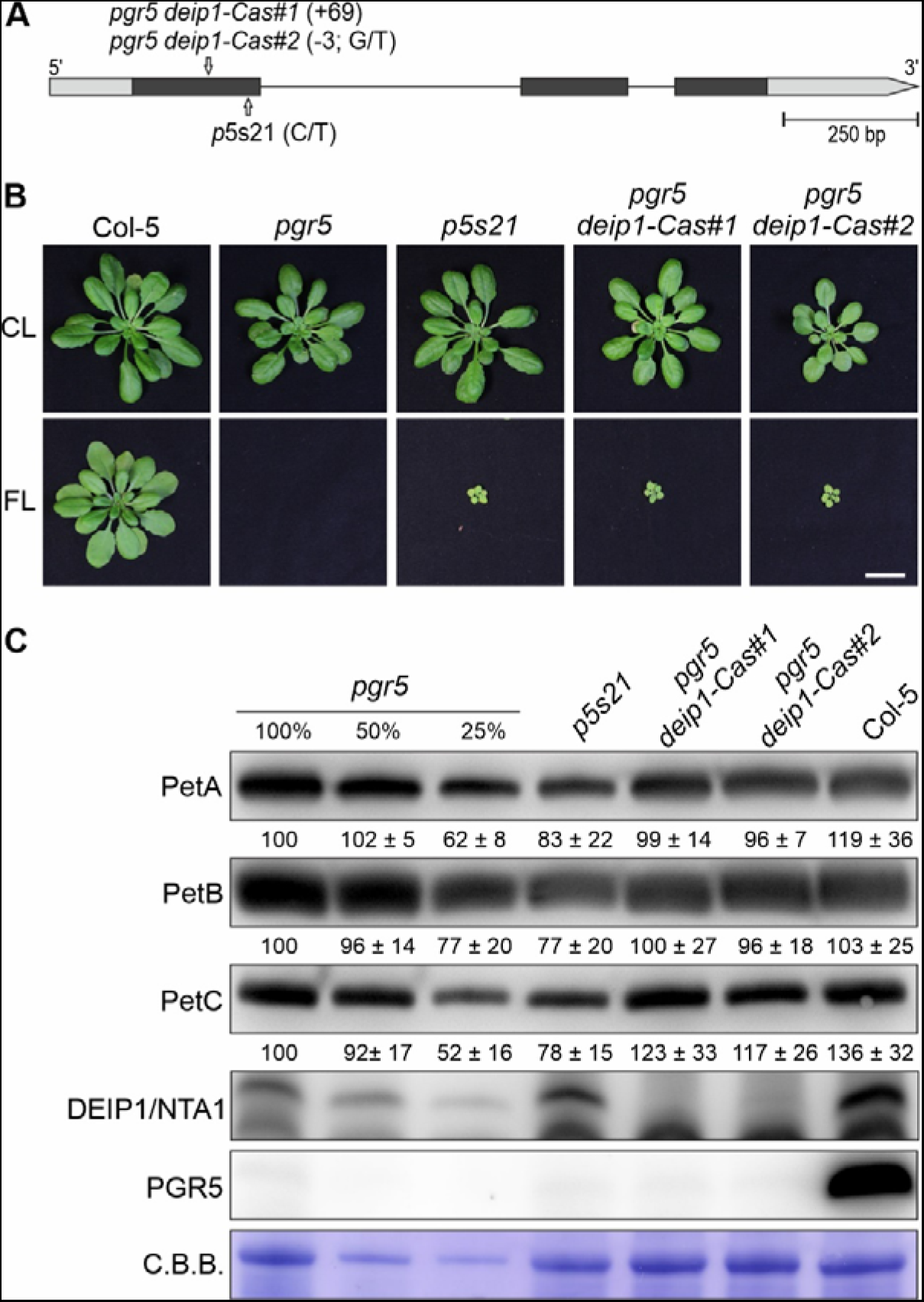
Growth and cyt *b*_6_*f* protein accumulation in adult plants lacking both DEIP1/NTA1 and PGR5. **(A)** Schematic representation of the *A. thaliana DEIP1*/*NTA1* coding sequence. The positions of the nucleotide insertions in the *pgr5 deip1-Cas#1* and *#2* mutants are shown, as well as the nucleotide substitution C/T in *p5s21*. The 5’- and 3’-UTR regions are shown as grey boxes, the exons in black, and the intron as a line. **(B)** Growth phenotype of WT (Col-5), *pgr5* and *pgr5 deip1/nta1* (*p5s21*, *pgr5 deip1-Cas#1* and *pgr5 deip1-Cas#2*) plants under control (5 weeks CL) and FL (5 weeks). Note that the *deip1*/*nta1* single mutant could not be obtained due to its lethality. Scale bar indicates 1 cm. **(C)** Aliquots of total leaf proteins were isolated from the CL-grown plants in (B), fractionated by SDS-PAGE under reducing conditions, transferred onto PVDF membranes, and immunodecorated with cyt *b*_6_*f*- (PetA, PetB, PetC), NTA1- or PGR5-specific antibodies. Coomassie Brilliant Blue (C.B.B.) staining of the membranes served as a loading control. Representative blots from three experiments are presented, as well as the average of the band quantifications (n=3) relative to *pgr5* (100%) ± SD.

Interestingly, we observed that at seedling stages (4-day-old plants, **Figure 5A**), the levels of cyt *b*_6_*f* in *pgr5 deip1-Cas#1* and *#2* were not recovered to the WT as in adult plants and remained lower than in *pgr5*, especially PetC, which was between 33% and 41% of *pgr5* (**Figure 5B**). Thus, the differences observed at the protein level resulted in slightly lower PSII activity in the double mutants compared to *pgr5*, without changes in NPQ (**Figure 5C**), which would explain the suppressor effect under FL due to increased PSI donor side limitation.

**Figure 5.**
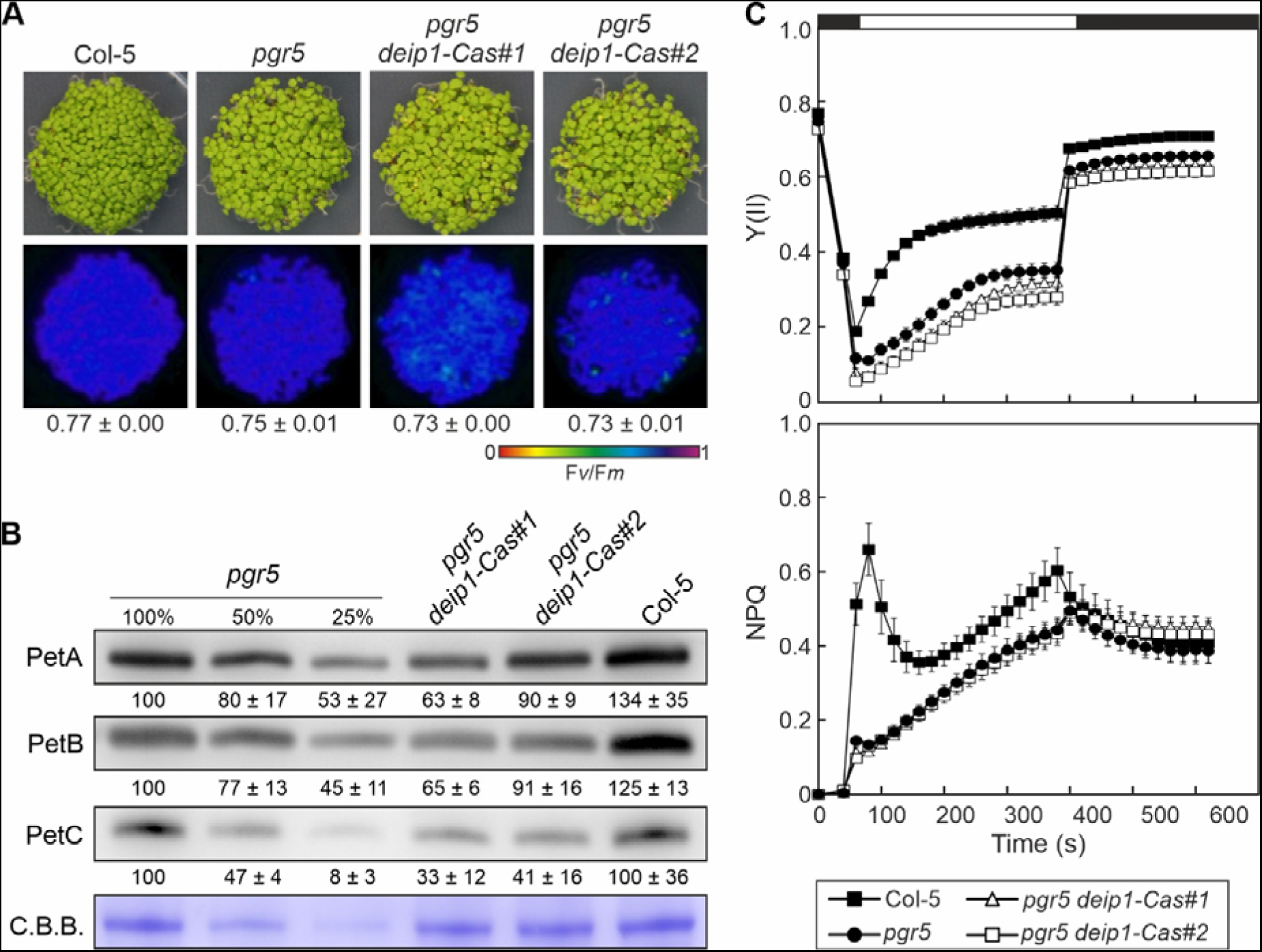
Cyt *b*_6_*f* protein accumulation and photosynthesis in seedlings lacking both DEIP1/NTA1 and PGR5. **(A)** Four-day-old WT (Col-5), single (*pgr5)* and double (*pgr5 deip1-Cas#1* and *pgr5 deip1- Cas#2*) mutant plants were grown on MS plates under long-day conditions. Below, false-color images show the F*v*/F*m* values for each plantlet line, according to the color scale at the bottom of the panel. For each line, the average of the values (n=3) ± standard deviation (SD) is shown. **(B)** Aliquots of total leaf proteins (adjusted to equal fresh weight) were isolated from the seedlings in (A), fractionated by SDS-PAGE under reducing conditions, transferred onto PVDF membranes, and immunodecorated with PetA-, PetB- or PetC-specific antibodies. Staining of the membranes with Coomassie brilliant blue (C.B.B.) served as loading control. Representative blots from three experiments are presented, as well as the average of the band quantifications (n=3) relative to *pgr5* (100%) ± SD. **(C)** PSII quantum yield (Y(II)) and non-photochemical quenching (NPQ) of dark-adapted seedlings in (**A**) during an induction-recovery curve (indicated by the white and black bars above) using the HEXAGON-IMAGING-PAM fluorimeter and applying 100 μmol photons m^−2^ s^−1^ of actinic light. Averages of at least 3 replicates ± SD are shown.

Taken together, it appears that DEIP1/NTA1 does not function as a simple assembly factor of the cyt *b*_6_*f* complex, since its absence can be fully compensated, at least in adult plants, and partially in very young plants, in the absence of PGR5. This suggests that one or more of the pleiotropic effects of the *pgr5* mutation on photosynthesis and thylakoid redox state render the function of DEIP1/NTA1 obsolete.

### At2g04360 is a chloroplast protein conserved in photosynthetic organisms

We found three suppressor lines, *p5s22*, *p5s23*/*24* and *p5s25*, with mutations in the *AT2G04360* gene (**Table 1**, **Figure 1** and **6A**). In silico analysis showed that the At2g04360 protein contains six transmembrane domains (**Figure 6B**) and is highly conserved in all photosynthetic organisms, including algae and cyanobacteria (**Figure 6B, C**). However, nothing is known about the function of At2g04360 in any of these organisms. Moreover, the encoded protein has a predicted chloroplast transit peptide (cTP), with the mature protein most likely starting at amino acid 66 according to TargetP 2.0 (**Figure 6B**). Indeed, At2g04360 was identified in the stromal lamellae of thylakoids in a previous proteomic analysis (Tomizioli et al., 2014). We experimentally confirmed the chloroplast location of At2g04360 by transient expression of an At2g04360-GFP fusion in isolated Arabidopsis protoplasts (**Figure 6D**).

**Figure 6.**
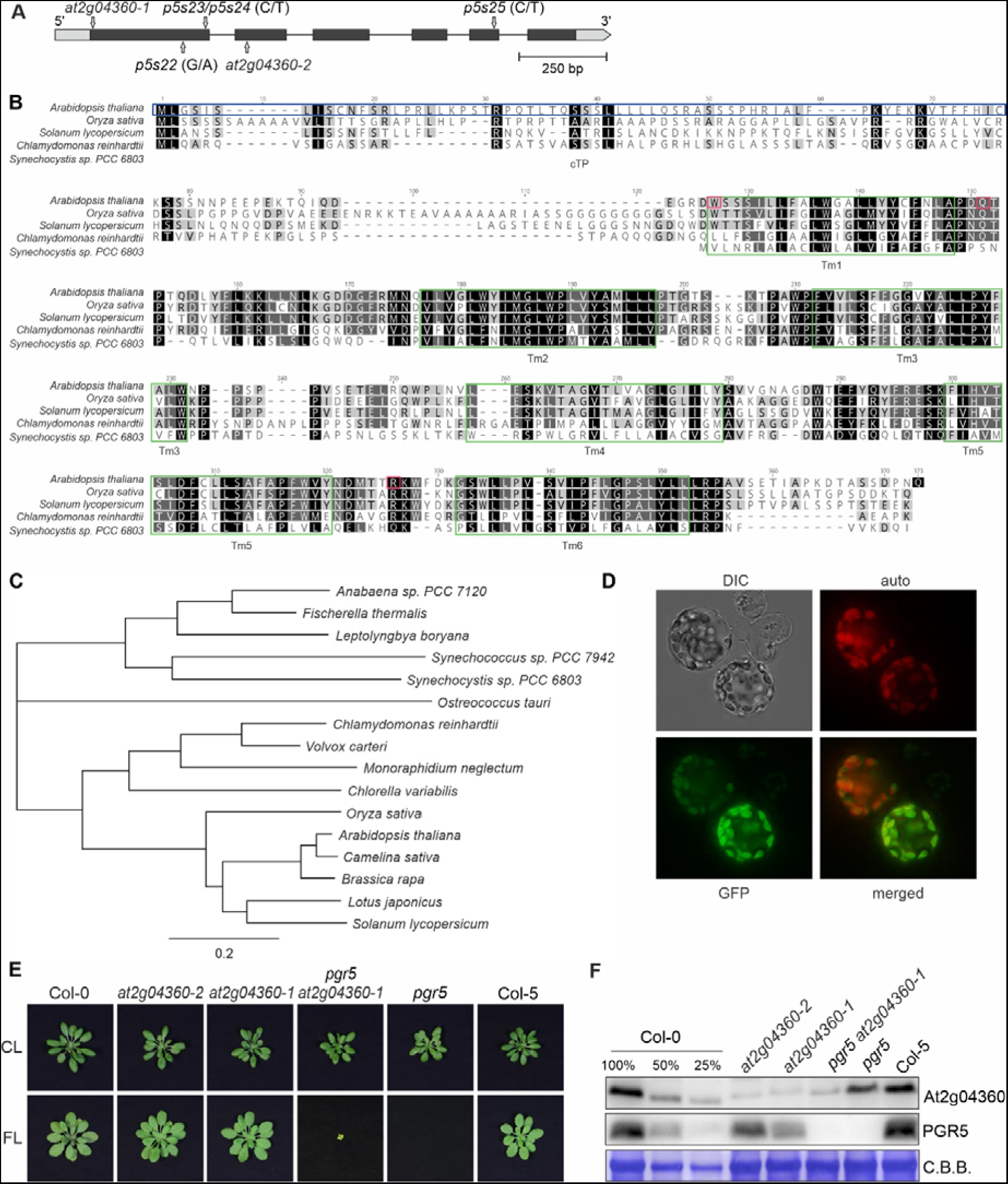
At2g04360 is a conserved chloroplast protein. **(A)** The *AT2G04360* gene: structure and T-DNA insertions in mutant lines. The position of the T-DNA in the *at2g04360-1* (from the GK collection) and *at2g04360-2* (from the Salk collection) is shown, as well as the nucleotide substitutions in *p5s22*, *p5s23/24* and *p5s25*. The 5’- and 3’-UTR regions are shown as gray boxes, the exons are shown in black, and the intron is shown as a line. **(B)** Multiple sequence alignment of At2g04360 proteins from different photosynthetic organisms. Instances of sequence identity/similarity in at least 40% of the sequences are highlighted by black/gray shading. The green rectangles indicate the transmembrane domains (Tm) of *A. thaliana* At2g04360 (Q29Q44). In the *A. thaliana* sequence, the blue rectangle indicates the predicted chloroplast transit peptide (cTP). The PFSC1-specific antibody was raised against the epitope comprising amino acids 65-81. The lines *p5s22*, *23/24* and *25* contain mutations leading to stop codons at the positions 88, 112 and 274 of *A. thaliana* At2g04360, respectively (magenta squares). **(C)** Phylogenetic tree using mature At2g04360 protein sequences (without cTPs) from different organisms, including plants, algae and cyanobacteria. **(D)** Subcellular location of At2g04360 determined by transient expression of At2g04360-GFP in Arabidopsis protoplasts. **(E)** Growth phenotype of WT (Col-0 and Col-5) and mutant (*at2g04360-2*, *at2g04360-1, pgr5, and pgr5 at2g04360-1*) plants under control (5 weeks of CL) and FL (5 weeks) conditions. Scale bar indicates 1 cm. **(F)** Aliquots of total leaf proteins were isolated from the CL-grown plants in (E), fractionated by SDS-PAGE under reducing conditions, transferred onto PVDF membranes, and immunodecorated with At2g04360- or PGR5-specific antibodies. Coomassie Brilliant Blue (C.B.B.) staining of the membranes served as a loading control.

To study the function of At2g04360, we identified two independent Arabidopsis T-DNA mutants with insertions at the beginning of the gene (*at2g04360-1*) and in the second exon (*at2g04360-2*) (**Figure 6A**). The two mutant lines grew similarly to WT under control and FL conditions (**Figure 6E**). However, it is worth noting that in both conditions and for both genotypes, we also found plants that were much smaller than WT. In addition, to prove that the mutation of *AT2G04360* really suppresses the lethal *pgr5* phenotype under FL, we generated the double mutant *pgr5 at2g04360-1* by crossing the two single mutants. Indeed, the double mutant was able to survive under FL conditions (**Figure 6E**), even up the flowering stage. Furthermore, the generation of an antibody against a peptide of At2g04360 allowed the detection of the protein in whole cell extracts, with a size of approximately 17-20 kDa (**Figure 6F**).

### At2g04360/PFSC1 is required at early developmental stages for cyt *b*_6_*f* accumulation and HL tolerance

To investigate the function of At2g04360, we switched our attention to seedlings to avoid the aforementioned heterogeneity observed in the growth of adult plants lacking At2g04360. At this stage of development stage, no differences in growth or F*v*/F*m* were observed between the different lines (**Figure 7A**). We then performed a proteomic analysis using pools of 4-day-old seedlings grown on MS plates (**Figure 7B**). 37 differentially expressed proteins (DEPs) were found in *at2g04360-1* compared to Col-0, considering those with at least two-fold change (FC) and less than 0.05 adjusted p-value (**Figure 7B**, **Supplemental Table S1, Supplemental Dataset 1**). Of these, 16 were up-regulated and 21 were down-regulated, most of which were located in plastids (**Figure 7C**). Interestingly, the levels of two cyt *b*_6_*f* subunits, PetA and PetB, were significantly altered in *at2g04360-1*, being about 30% of the WT. The same level was observed for PetC, although the change was not statistically significant. The changes in cyt *b*_6_*f* accumulation were specific regarding photosynthesis, as no changes in other photosynthetic proteins were found (**Supplemental Table S1**). The results of the proteomics analysis were corroborated by immunoblot analysis, which also included the *at2g04360-2* and *pgr5 at2g04360-1* lines (**Figure 7D**). All three *pfsc1* lines showed reduced levels of PetA, PetB and PetC, while PsbE/F and PsaD1 remained similar to WT (**Figure 7D**). These results suggest that At2g04360 is required for the accumulation of cyt *b*_6_*f*, particularly in early developmental stages, but unlike DEIP1/NTA1, independent of PGR5, since lower levels of this complex were also found in *pgr5 at2g04360-1*. Because of the altered cyt *b*_6_*f* levels, we named the mutation “*<u>p</u>gr<u>5 s</u>uppressor with altered <u>c</u>yt b_6_f <u>1</u>*” or *pfsc1*, whereby the “*f*” stands for “five”, and the *AT2G04360* gene *PFSC1*.

**Figure 7.**
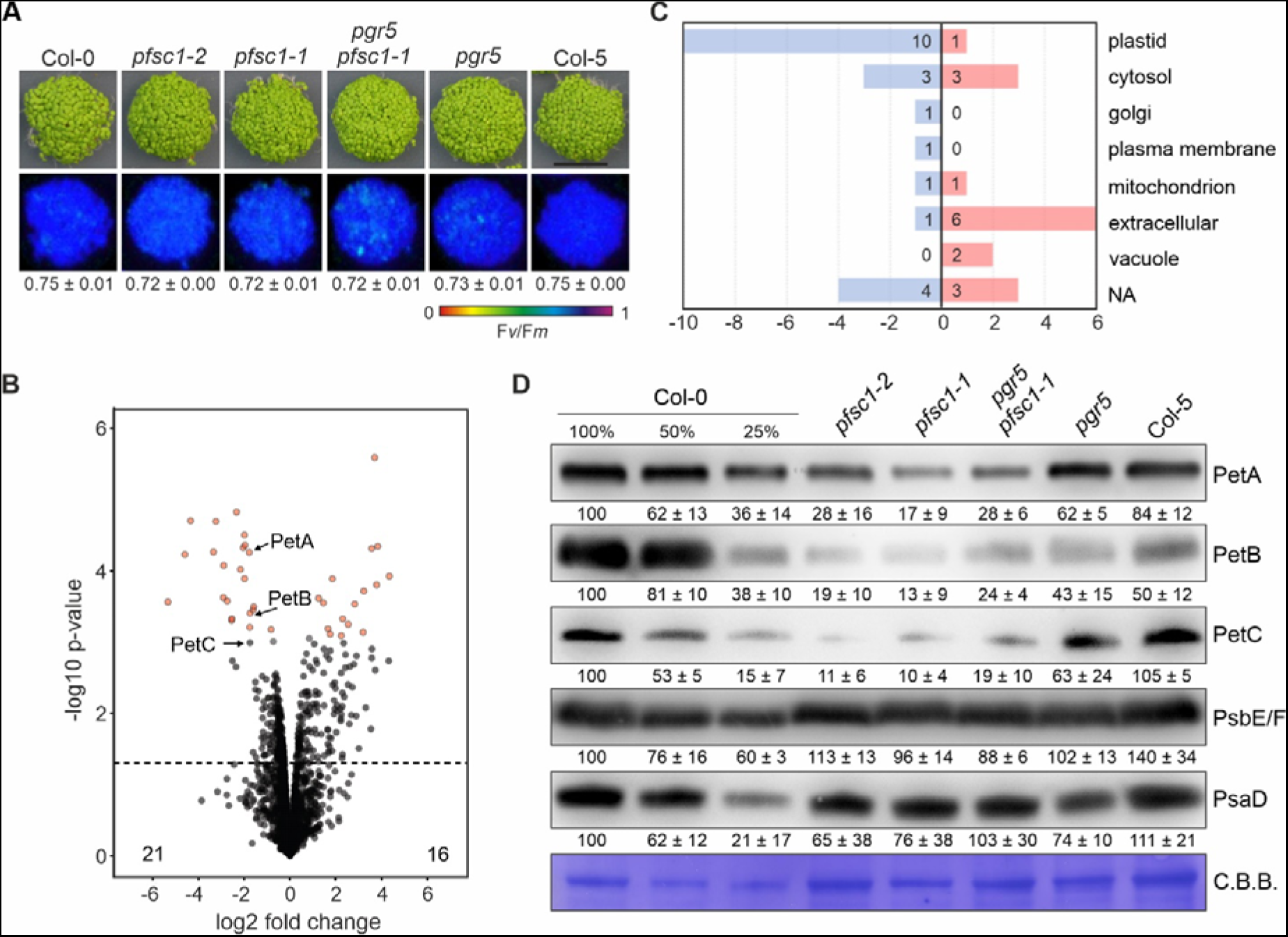
Maximum PSII quantum yield and cyt b_6_*f* accumulation in seedlings lacking At2g04360/PFSC1. **(A)** Four-day-old WT (Col-0 and Col-5), single (*pgr5, at2g04360-1/pfsc1-1* and −2) and double (*pgr5 at2g04360-1/pfsc1-1*) mutant plants were grown on MS plates under long-day conditions. Below, false-color images show the F*v*/F*m* values for each plantlet line, according to the color scale at the bottom of the panel. For each line, the average of the values (n=3) ± (SD) is shown. **(B)** Volcano plot of differentially (*at2g04360-1*/*pfsc1-1* vs. Col-0) expressed proteins (DEPs) as in (A). Each point indicates a different protein, ranked according to P-value (y-axis, −log10 of P-values) and relative abundance ratio (x-axis, log2 Fold Change *pfsc1-1* Col-0). DEPs (log2 FC > |1|, FDR < 0.05) are labelled in red. The dashed line indicates a negative log10 P-value of 1.3. **(C)** Subcellular distribution of the DEPs shown in (B). Subcellular location was determined using the SUBA4 database. **(D)** Aliquots of total leaf proteins (adjusted to equal fresh weight) were isolated from seedlings as shown in (A), fractionated by SDS-PAGE under reducing conditions, transferred onto PVDF membranes, and immunodecorated with PetA-, PetB-, PetC-, PsbE/F- and PsaD-specific antibodies. Staining of the membranes with Coomassie Brilliant Blue (C.B.B.) served as loading control. Representative blots from three experiments are presented, as well as the average of the band quantifications (n=3) relative to Col-0 (100%) ± SD.

The analysis of the photosynthetic performance (**Figure 8A**) showed that seedlings lacking PFSC1 had very low electron transport rates from PSII (ETR(II)), despite F*v*/F*m* values higher than 0.7 (**Figure 7A**). Furthermore, they were unable to induce transient NPQ, which was even lower than in *pgr5*, and showed a strongly reduced plastoquinone pool during illumination (1-qL close to 1). These results are consistent with the defective cyt *b*_6_*f* accumulation observed in the absence of PFSC1, as these plants will have a bottleneck of electrons on plastoquinones, as well as impaired acidification of the lumen. Additionally, the lower electron transport in *pgr5 pfsc1-1* compared to *pgr5* at the seedling stage (**Figure 8A**), may explain the survival phenotype of these plants under FL conditions.

**Figure 8.**
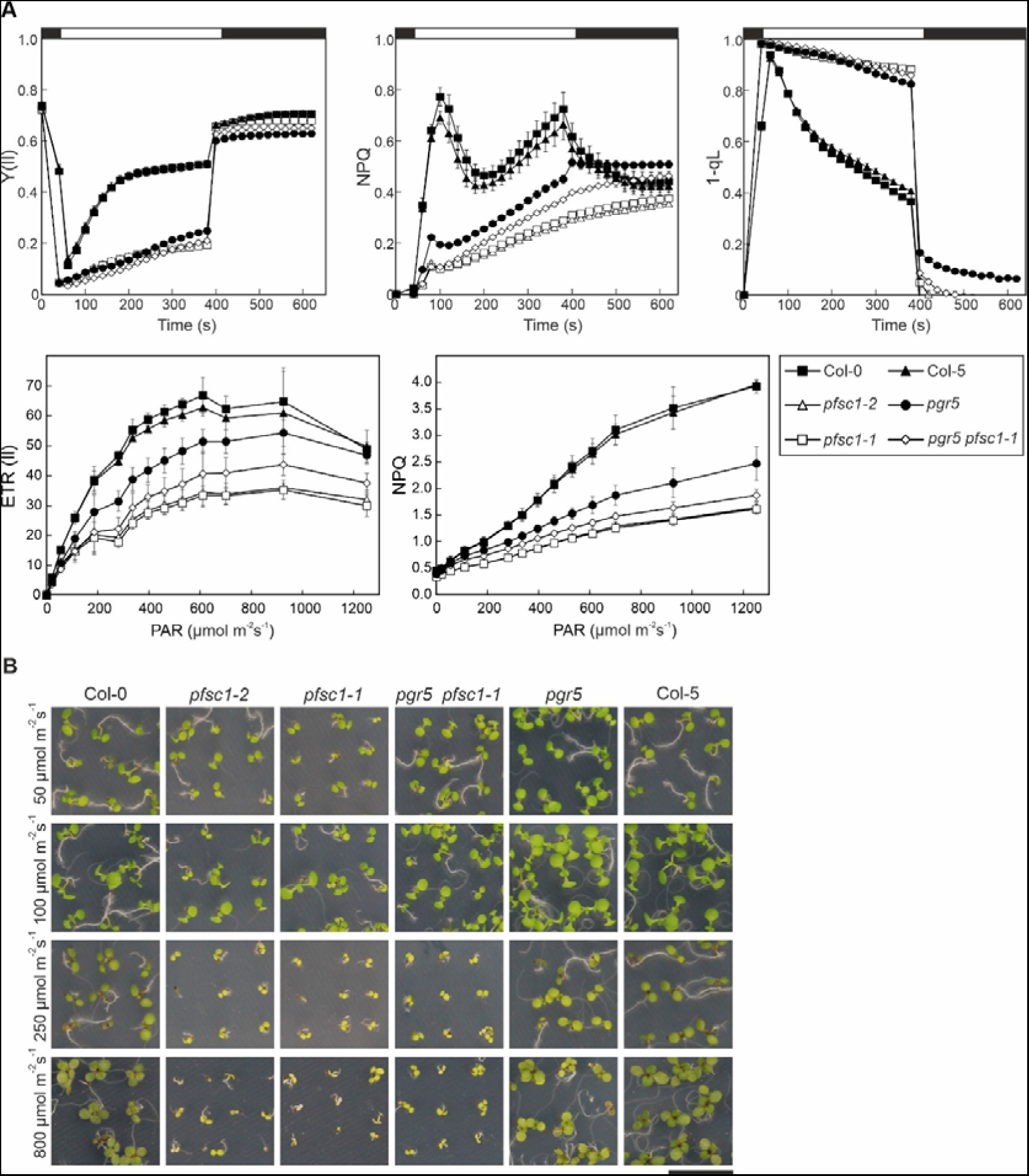
PFSC1 is required for proper photosynthesis and high light tolerance in seedlings. **(A)** Measurement of photosynthetic parameters. PSII quantum yield (Y(II)), non-photochemical quenching (NPQ) and the redox state of the plastoquinone pool (1-qL) (upper three panels) were determined in 4-day-old WT (Col-0 and Col-5), single (*pgr5, pfsc1-1* and *-2*) and double (*pgr5 pfsc1-1*) mutant plants grown on MS plates under long-day conditions, during an induction-recovery curve (indicated by the white and black bars above) using the HEXAGON-IMAGING-PAM fluorimeter and applying 100 μmol photons m^−2^ s^−1^ of actinic light. Plants were previously dark adapted for 30 min. Electron transport rates from PSII (ETR(II)) and NPQ (lower two panels) were measured at increasing photosynthetically active radiation (PAR) using the IMAGING-PAM fluorimeter. Averages of at least 3 replicates ± SD are shown. **(B)** Growth phenotype of the same genotypes as in A grown on MS plates under long-day conditions and different light intensities for seven days. Scale bar indicates 1 cm.

Correspondingly, young plants lacking PFSC1 were very sensitive to high light (**Figure 8B**). The mutants died after the cotyledon stage at 800 and even at 250 µmol photons m^−2^ s^−1^. In contrast, at lower light intensities (50-100 µmol photons m^−2^ s^−1^), these plants were able to grow like the WT (**Figure 8B**). Autoshading and its absence is therefore the most likely explanation for the aforementioned observation that among *pfsc1* plants, a certain proportion of plants were found that were much smaller than WT.

## DISCUSSION

In our screen, we identified twelve proteins whose mutation can restore the viability of *pgr5* plants under FL. This portfolio of proteins allows us to confirm previous models, based on photosynthesis measurements, of how the harmful effects of the *pgr5* mutation on PSI can be compensated (Yamamoto and Shikanai, 2019). The screen uncovers novel facets of the relationship of PGR5- and NDH-dependent CEF pathway. Finally, the screen also identifies proteins with functions in the accumulation or protection of cyt *b*_6_*f*.

### Suppressors of *pgr5* are affected in photosynthesis-related proteins

As the first four mutations identified in the suppressor screen all affected PSII and caused markedly decreased F*v*/F*m*, we subsequently selected suppressor mutants with little or no effect on F*v*/F*m* to uncover the effect of other chloroplast functions beyond PSII on PGR5-dependent CEF. Nevertheless, we found mutations in proteins associated with PSII (PPL1 and Psb33, see **Table 1**) that do not affect F*v*/F*m*, and it is likely that these mutations cause perturbations in the LEF at early developmental stages. Other *pgr5* suppressor mutations were found to affect proteins already known to be involved in cyt *b*_6_*f* assembly (CcdA) (Page et al., 2004), copper transport relevant to plastocyanin accumulation (PAA1) (Shikanai et al., 2003), and cFBP1 (Rojas-Gonzalez et al., 2015) and one of its regulators (ACHT2) (Yokochi et al., 2021). The remaining two proteins targeted by suppressor mutations have functions associated with cyt *b*_6_*f*. PFSC1, with previously unknown functions, controls cyt *b*_6_*f* accumulation at early developmental stages. DEIP1/NTA1, previously identified as a cyt *b*_6_*f* assembly factor (Sandoval-Ibanez et al., 2022; Li et al., 2023), functionally interacts with PGR5 in a completely unexpected manner, suggesting a novel link between PGR5 function and cyt *b*_6_*f* accumulation. As a consequence, the screen uncovered mutations exclusively in photosynthetic components, suggesting that there are probably no additional mechanisms other than photosynthetic regulation that can compensate for the absence of PGR5.

Equally interesting is the question of why some mutations were not found in the screen, such as PSI subunits, the *pgr1* mutation already described to antagonize the effects of *pgr5* (Yamamoto and Shikanai, 2019), and mutation that cause high NPQ to compensate for the defect in transient NPQ induction of the *pgr5* mutant. We addressed this question by introducing a mutation of *PSAD1*, which affects the accumulation of the D subunit of PSI (Ihnatowicz et al., 2004), and the *pgr1* mutation into the *pgr5* background. Both mutations could not restore viability under FL in the *pgr5 psad1* and *pgr5 pgr1* double mutants, although both mutations also caused a strong increase in PSI donor side limitation under HL in adult plants, as did the *pam68* mutation, which could rescue FL lethality. However, the corresponding double mutants did not display the same increased PSI donor-side limitations as *psad1* and *pgr1* did, although *pgr5 pgr1* showed a moderate increase in PSII donor side limitation, but not to the same extent as for example *p5s1*, *14* and *15* and *pgr5 pam68* (see **Figure 3**). Therefore, there may be a threshold level of PSII donor side limitation that needs to be reached to achieve viability under FL.

As transient NPQ induction is reduced in *pgr5*, one might also expect to identify mutations that restore NPQ in the screen, given the protective role of NPQ in preventing ROS formation and photoinhibition. However, we have previously shown that this is not the case by introducing two high NPQ mutations into the *pgr5* background (Naranjo et al., 2021). Neither the *cgl160* mutation, which is defective in the thylakoid ATP synthase and therefore has a lower rate of proton consumption, nor the inactivation of NTRC could rescue the lethal phenotype of *pgr5*, although the *cgl160* mutation could at least partially restore the transient NPQ induction (Naranjo et al., 2021).

### Mechanisms of *pgr5* suppression

Previous studies have described that the addition of DCMU, which inhibits PSII (Suorsa et al., 2012), or the mutants Δ*5 pgr5* (Suorsa et al., 2016) and *pgr1 pgr5* (Yamamoto and Shikanai, 2019), protect PSI and restore photosynthetic parameters in the *pgr5* background. Furthermore, the lack of PGR5 can also be compensated by increasing the electron sink capacity from PSI, for example by adding methyl viologen, which directly accepts electrons from PSI (Munekage et al., 2002), or by overexpressing *Physcomitrella patens* flavodiiron protein genes in *pgr5* (Yamamoto and Shikanai, 2019). Our analysis of the photosynthetic electron flow of the suppressor lines indicated that this concept easily explains the recovery of suppressor lines under FL, namely a decrease in electron flow to PSI and/or an increase from PSI to its acceptors. Since our screen was based on the recovery of lethality of *pgr5* seedlings, the effect of the second-site mutations on PSI donor and acceptor side limitation must occur at early stages of plant development. Nevertheless, we were able to observe these effects on the PSI donor and acceptor sides in adult plants with impaired PSII (*p5s1* and *pgr5 pam68*), cyt *b*_6_*f* (*p5s14*) or PC (*p5s15*) (see **Figure 3**). In the four lines with mutations associated with cFBP1 (*p5s18/19* and *p5s20*), DEIP1/NTA1 (*p5s21*) or At2g04360 (*p5s23*), however, the effects were no longer detectable in adult plants. However, when we examined early developmental stages of suppressor lines carrying mutations in DEIP1/NTA1 (*p5s21*) (see **Figure 5**) or PFSC1 (*p5s23*) (see **Figure 8**), we could confirm that they were indeed altered in their photosynthetic electron flow to PSI.

We found a suppressor line in which electron flow from PSI to acceptors may be increased. Line *p5s20* is defective in ACHT2, a thioredoxin-like protein that down-regulates the Calvin-Benson cycle by oxidative inactivation of the cFBP1 (Yokochi et al., 2021). Thus, a deficiency of ACHT2 could result in a more active CO_2_ fixation, providing an increased sink capacity for PSI-derived electrons. This hypothesis needs to be tested in follow-up analyses. In addition, it has been previously described that plants with an inactive *CFBP1* gene have an overaccumulation of the NDH complex, resulting in hyperactivity of NDH-CEF (Livingston et al., 2010). Furthermore, the *pgr5 cfbp1* mutation was described to have less light-saturated LEF compared to *cfbp1*, while the *cfbp1 crr2* double mutant with a defective NDH complex was severely impaired in photosynthesis and growth, leading to the conclusion that the *cfbp1* mutation imposes a demand for more ATP production, which was met by increased NDH-dependent CEF that could not be replaced by PGR5-CEF (Livingston et al., 2010). Our study has now shown that the severe growth retardation of plants that lack cFBP1 is suppressed by the *pgr5* mutation (see **Figure 2**), shedding new light on the findings of Livingston et al. (2010). Our results provide an indirect support for the paradigm that PGR5 is primarily involved in CEF and show that the capacity of PGR5-CEF is physiologically significant. The most parsimonious explanation for the suppression of the *cfbp1-1*-related growth phenotype by *pgr5* is that the depletion of PGR5-CEF rebalances the overall level of CEF to WT-like levels. However, this also implies that the magnitude of PGR5-dependent CEF must be in the range of the hyper-activity of NDH-dependent CEF in *cfbp1-1* and is not insignificant, as concluded by Livingston et al. (2010). The real question here is why NDH-CEF and not PGR5-CEF is up-regulated in *cfbp1-1*? One possibility is that PGR5-CEF cannot be further enhanced, but other studies have shown that a defect in PSI can indeed result in enhanced PGR5-CEF (DalCorso et al., 2008). Another possibility is that under certain conditions, enhancing NDH-CEF is safer than enhancing PGR5-CEF, and this possibility is also discussed below in the context of DEIP1/NTA1. Indeed, previous studies suggest that enhanced PGR5 levels (Okegawa et al., 2007; Long et al., 2008) or the absence of its regulators (Rühle et al., 2021) can have detrimental effects on photosynthesis.

### Novel cyt *b*_6_*f* related functions for DEIP1/NTA1 and PFSC1

Our *pgr5* suppressor screen also found mutants with altered LEF due to impaired cyt *b*_6_*f* accumulation in early developmental stages. These include a mutation in CcdA in the suppressor line *p5s14*, which is involved in cyt *b*_6_*f* assembly (Page et al., 2004) (see **Table 1** and **Figure 1**). The effects of the *pgr1* mutation in the *PETC* gene on LEF were not sufficient to suppress the lethality of the *pgr5* mutation (see above). We also identified mutations in another bona fide cyt *b*_6_*f* biogenesis factor, DEIP1/NTA1, in the suppressor line *p5s21*. The DEIP1/NTA1 protein has been shown to interact with cyt *b*_6_*f* in two previous studies, but with some discrepancies: in one study it was found to interact with PetA and PetB (Sandoval-Ibanez et al., 2022), and in the other with PetB, PetD, PetG, and PetN (Li et al., 2023). This, together with the observation that plants lacking this protein are unable to accumulate the complex and are therefore seedling lethal, led to the conclusion that DEIP1/NTA1 is essential for cyt *b*_6_*f* biogenesis (Sandoval-Ibanez et al., 2022; Li et al., 2023). However, the finding that plants lacking DEIP1/NTA1 and PGR5 are viable and only show moderate decreases in the levels of representative cyt *b*_6_*f* subunits at early stages of development stages, contradicts the idea that DEIP1/NTA1 is essential for cyt *b*_6_*f* biogenesis. It appears that DEIP1/NTA1 is critical for cyt *b*_6_*f* accumulation only when PGR5 is present, suggesting that PGR5 has a damaging effect on the cyt *b*_6_*f* complex and that DEIP1/NTA1 protects the complex from this damaging effect. The mechanisms by which PGR5 exerts this deleterious effect remains elusive, but it is not unknown that PGR5 can exert deleterious effects, as overexpression of PGR5 or removal of its regulators PGRL1 and PGRL2 has been shown to be associated with deleterious effects on photosynthesis and growth (Okegawa et al., 2007; Long et al., 2008; Rühle et al., 2021). A direct role of PGR5 in this destabilization of cyt *b*_6_*f* cannot be excluded either, since PGR5 actually interacts with the Cyt *b*_6_ protein in split-ubiquitin assays.

Three independent suppressor lines (*p5s22*, *23/24* and *25*) were mutated in the *PFSC1* gene. The PFSC1 protein is required for the accumulation of cyt *b*_6_*f* in seedlings (see **Figure 7**), and both the sensitivity of *pfsc1* seedlings to high light (see **Figure 8**) and the suppression of the *pgr5* lethality under FL conditions can be attributed to the significant reduction in cyt *b*_6_*f* accumulation at this stage of development. However, this appears to occur in a different manner than DEIP1/NTA1, because unlike DEIP1/NTA1, PFSC1 functions independently of PGR5, as shown by the similar phenotypes observed in the *pgr5 pfsc* and *pfsc* plants with respect to cyt *b*_6_*f* protein accumulation (see **Figure 7**). Furthermore, PFSC1 is present in all photosynthetic organisms, including cyanobacteria (see **Figure 6**), whereas DEIP1/NTA1 is exclusive to plants and green algae (Sandoval-Ibanez et al., 2022; Li et al., 2023). Interestingly, seedling plants lacking PFSC1 express only 10-28% of the cyt *b*_6_*f* subunits tested, but are able to grow similarly to WT plants and have WT-like Fv/Fm values under low light conditions (see **Figure 7**).

### Outlook

The *pgr5* screen was able to identify those steps in photosynthesis that are affected by PGR5 deficiency, and that need to be mutated to compensate for the lack of PGR5. Thus, in addition to supporting the concept of Yamamoto and Shikanai (2019) that PGR5-dependent CEF protects PSI under FL on the donor and acceptor side, and the possibility of identifying novel factors required for the biogenesis of photosynthetic proteins involved in LEF (such as PFSC1), the screen has uncovered that PGR5 can have a deleterious effect on photosynthesis. This detrimental effect becomes apparent when cFBP1 is inactivated, resulting in a down-regulation of PGR5 accumulation as described before (Livingston et al., 2010) and in our study (see **Figure 2B**). Only complete inactivation of PGR5-dependent CEF by introducing the *pgr5* mutation in the *cfpb1* background restores WT-like growth. This behavior is similar to that observed in the *pgr5 ntrc* mutant, where complete removal of PGR5 results in recovery of growth, whereas the *ntrc* mutant exhibits severe growth retardation and a decrease in PGR5 levels (Naranjo et al., 2021). This suggests that some mutations create a situation where PGR5 expression must be downregulated to prevent its deleterious effects. Moreover, the deleterious effect of PGR5 on the accumulation of the cyt *b*_6_*f* complex appears to have necessitated the evolution of the DEIP1/NTA1 protein for protection, possibly indicating that this deleterious effect of PGR5 is specific to eukaryotes, since DEIP1/NTA1 is not present in cyanobacteria.

The *pgr5* screen was not designed to identify direct regulators of PGR5. For this purpose, the concept of suppressing lethality under FL needs to be extended. For example, screening for second site mutations that suppress lethality due to absence of PGRL1 is one way to further disentangle the mechanisms and regulation of PGR5-dependent processes in photosynthesis.

## METHODS

### Plant material

In this study, *Arabidopsis thaliana* Col-0 and Col-5 were used as WT controls. The point mutation lines *pgr5* (Munekage et al., 2002), *pgr1* (Munekage et al., 2001) and *pgr5 pgr1* (Yamamoto and Shikanai, 2019), as well as the T-DNA lines *cfbp1-1* (Rojas-Gonzalez et al., 2015), *pam68* (Armbruster et al., 2010), *psad1* (Ihnatowicz et al., 2004), and *pgrl1ab* (DalCorso et al., 2008) have been described previously. The double mutants *pgr5 pam68*, *pgr5 psad1* and *pgr5 pgr1* were generated by manual crossing of the corresponding single mutants and genotyped using the primers listed in **Supplemental Table S2**. The single mutant *acht2-Cas* and the double mutants *pgr5 acht2-Cas*, *pgr5 deip1-Cas#1* and *pgr5 deip1-Cas#2* were obtained by the introduction of a CRISPR/Cas9 construct specific to the respective target genes *AT4G29670* and *AT2G27290* into the Col-0 or *pgr5* background as described (Penzler et al., 2022). A vector with an oocyte-specific promoter, pHEE401-E (Wang et al., 2015), was combined with the specific guide RNA (gRNA) for *AT4G29670* and *AT2G27290* (**Supplemental Table S2**). Transformation of Col-0 or *pgr5* was accomplished via *Agrobacterium tumefaciens* GV3101 using the floral dipping method (Clough and Bent, 1998). The selection of transgenic lines was carried out on plates containing Murashige and Skoog (MS) salt medium (1x), 25 μg mL^−1^ hygromycin, 1% [w/v] plant agar, and 1% [w/v] sucrose, in the first generation after transformation, T_1_. The genes were then sequenced using the primers in **Supplemental Table S2** to identify possible mutations. Lines in which the targets (*ACHT2* or *DEIP1/NTA1*) were affected by a mutation were propagated and selected in the T_2_ generation to obtain stable homozygous mutant lines. The T-DNA mutants *at2g04360-1*/*pfsc1-1* (GK-108A10) and *at2g04360-1*/*pfsc1-2* (SALK_206798) were obtained from the Nottingham Arabidopsis Stock Centre (NASC) and selected by genotyping using the primers in **Supplemental Table S2**. The double mutant *pgr5 pfsc1-1* was generated by manual crossing of the respective single parental lines.

### Plant growth conditions

Seeds from mutant and WT plants were sown in potting soil and stratified at 4°C for three days. They were then cultivated in climate chambers under various light conditions and day lengths: Long day (LD), 16 h light (100 µmol photons m^−2^ s^−1^) / 8 h darkness; Mild high light (mHL), 16 h light (250 µmol photons m^−2^ s^−1^) / 8 h darkness; High light (HL), 16 h light (800 µmol photons m^−2^ s^−1^) / 8 h darkness; Control light (CL), 12 h light (100 µmol photons m^−2^ s^−1^) / 12 h darkness; and Fluctuating light (FL), 12 h light / 12 h darkness, with cycles of 5 min at 50 μmol photons m^−2^ s^−1^ and 1 min at 500 μmol photons m^−2^ s^−1^ during the light period. The temperature was maintained at 22°C during the day and 20°C during the night, with a constant relative humidity of 60% under all conditions. Fertilizer was applied following the manufacturer’s recommendations (Osmocote Plus; Scotts, Nordhorn, Germany). For seedling experiments, seed sterilization was conducted as follows. Seeds were immersed in a solution containing 70% [v/v] EtOH and 0.01% [v/v] Tween-20 for 20 min, followed by immersion in 90% [v/v] EtOH for 10 min, all with continuous shaking. The sterilized seeds were then air-dried on sterile Whatman filter paper under aseptic conditions. The growth medium for the experiments consisted of 1% [w/v] MS medium (PhytoTechnology Laboratories, LLC™, USA) and 0.8% [w/v] agar. The sterilized seeds were then transferred to the prepared growth medium in Petri dishes and stored in the dark at 4°C for 48 h to undergo stratification. Subsequently, the MS plates were transferred to the specified growth conditions as detailed above.

### EMS mutagenesis and screening

Approximately 500-600 mg of freshly harvested seeds, corresponding to approximately 20,000 M_0_ seeds of the *pgr5* mutant, were subjected to mutagenesis as follows. The seeds were placed in a 50 ml tube containing 35 ml of H_2_O and 30 mM EMS (Sigma-Aldrich, St. Louis, USA). The EMS was dissolved before by shaking for 5 mins. Subsequently, the seeds were incubated in the EMS solution for 15 h with continuous shaking. After the incubation period, the seeds were rigorously washed 30 times with 50 ml of H_2_O to remove any residual EMS. The EMS solution was then neutralized by treatment with 0.1 M NaOH. The mutagenized seeds (M_1_) were air-dried on Whatman filter paper overnight and then sown on 40 trays containing soil in the greenhouse. Each tray represents one batch and contains approximately 500 M_1_ seeds. The seeds harvested from each batch (M_2_) were later used for selection under FL growth conditions. To saturate the screen, 273,000 M_2_ plants were screened from 20,000 M_1_ individuals (Maple and Moller, 2007). From the survivors, those with F*v*/F*m* values higher than 0.7 were selected. The *pgr5* suppressor candidates of interest were backcrossed with the parental line *pgr5* to remove additional mutations and selected again under FL in the second generation after crossing (F_2_).

### Whole-genome resequencing and data analysis

DNA for whole-genome resequencing was obtained from the *pgr5* suppressor candidates of interest in the F_2_ generation and after 5 weeks of selection under FL. As previously described (Penzler et al., 2022), leaves from a pool of 50-60 plants were ground in liquid nitrogen and incubated in lysis buffer (0.4 M sucrose, 10 mM Tris-HCl pH 7.0, 1% [v/v] β-ME, 1% [v/v] Triton) on ice for 15 min. The lysate was then filtered and centrifuged for 15 min at 1200 g and 4°C, and the pellet was resuspended in 1 mL of lysis buffer and centrifuged again for 15 min (600 g, 4°C). DNA was isolated from the supernatant using the DNeasy Plant Mini Kit (Qiagen, Hilden, Germany). Two µg of DNA were used to prepare 350-bp insert libraries for 150-bp paired-end sequencing (Novogene Biotech, Beijing, China) on an Illumina HiSeq 2500 system (Illumina, San Diego, USA) using standard Illumina protocols. The sequencing depth was at least 7 G of raw data per sample, which corresponds to higher than 50-fold coverage of the *A. thaliana* genome. After grooming the FASTQ files, adapters were removed using Trimmomatic (Bolger et al., 2014), reads were mapped to the TAIR10 annotation using BWA (Li and Durbin, 2009) with the parameters ‘mem -t 4 -k 32 -M’, and duplicates were removed using SAMtools (Li et al., 2009) with the rmdup tool. Single nucleotide polymorphisms (SNPs) were identified using SAMtools (Li et al., 2009) with the following parameters: ‘mpileup -m 2 -F 0.002 -d 1000’. Only SNPs that were supported by more than 8 reads and with mapping quality >20 were retained. To identify the SNPs specific for the candidate *pgr5* suppressors (*p5sX*), the SNPs between *p5sX* and *pgr5* were compared. The resulting *p5sX*-specific SNP list was submitted to the web application CandiSNP (Etherington et al., 2014), which generates SNP density plots. The output list of CandiSNP was screened for non-synonymous amino acid changes and the G/C to A/T transitions that were likely to be caused by EMS, with a particular focus on the chromosome with the highest density of SNPs with an allele frequency of >0.75.

### Chlorophyll fluorescence and P700 measurements in adult plants

To assess the photosynthetic performance in vivo, simultaneous measurements of chlorophyll *a* fluorescence and P700 absorbance changes were performed using a Dual-PAM-100 spectrophotometer (Walz, Effeltrich, Germany). The measurements were conducted on single attached leaves, dark adapted for 30 min, using a specific program that mimics the FL growth conditions used in the screen (**Supplemental Figure S2**). The maximum fluorescence (F*m*) and minimum fluorescence (F*_0_*), as well as the maximum absorption of PSI (P*m*), were determined by applying a saturation pulse (SP) with an intensity of 8,000 µmol photons m^−2^ s^−1^ for 0.3 sec. After 40 sec of darkness, the leaves were illuminated with 50 µmol photons m^−2^ s^−1^ of actinic light (low light, LL) for 5 min. Subsequently, the actinic light was switched to 500 µmol photons m^−2^ s^−1^ for 1 min (high light, HL). The LL / HL periods were repeated 5 and 4 times, respectively, for a total of 29 min and then, followed by a recovery phase of 8 min in darkness, while photosynthetic activity was continuously monitored. Saturation pulses were applied every 20 sec during the LL phase, every 15 sec during the HL phase, and every 20 sec during the recovery phase to determine the different parameters, which were then calculated by the DUAL-PAM software using the previously described equations (Klughammer and Schreiber, 2008b, a). The data were presented as the mean of three replicates.

### Chlorophyll fluorescence measurements in seedlings

The photosynthetic activity of PSII was determined in seedlings growing in MS plates by measuring the chlorophyll *a* fluorescence using the HEXAGON-IMAGING-PAM system (Walz, Effeltrich, Germany). The seedlings were dark-adapted for 30 min before the measurements. F*m* and F*_0_*were determined applying a SP with an intensity of 8,000 µmol photons m^−2^ s^−1^ for 0.3 sec. After 40 sec of darkness, a photosynthetic induction period of 360 sec with 100 µmol photons m^−2^ s^−1^ of actinic light took place, followed by a recovery phase of 180 sec of darkness. Yield measurements were recorded by applying saturation pulses every 20 sec, and the different parameters were then calculated by the IMAGING-PAM software using the equations described previously (Klughammer and Schreiber, 2008b, a). Light curves (LC) were obtained using an IMAGING-PAM spectrophotometer (Walz, Effeltrich, Germany) by applying stepwise increasing actinic light intensities every 3 min, and saturation pulses were applied at the end of each step to calculate photosynthetic parameters as described above.

### Protein extraction and immunoblot analysis

Rosette leaves or seedlings were immediately frozen in liquid nitrogen after collection and ground using a Tissuelyser II homogenizer (Qiagen, Venlo, Netherlands). The obtained powder was resuspended in 1 mL of 2x Tricine buffer per each 100 mg fresh weight of plant material. This buffer consisted of 8% [w/v] SDS, 24% [w/v] glycerol, 15 mM DTT and 100 mM Tris/HCl pH 6.8. After homogenization, the samples were heated at 70°C for 5 min and centrifuged at 13,000 g for 10 min, retaining the supernatant. The volume of solubilized proteins equivalent to 1 mg of fresh weight was loaded onto 10% [w/v] tricine-SDS polyacrylamide gels. An exception was the detection of PGR5 and NTA1 proteins, for which 3 and 30 mg fresh weight, respectively, were used. The separated protein samples were transferred to PVDF membranes (Immobilon-P; Millipore, Burlington, MA, USA) using the Trans-Blot Turbo system (Bio-Rad, Hercules, CA, USA). The PVDF membranes were blocked for 1 h with 5% [w/v] milk in Tris-buffered saline with Tween 20 (TBS-T; 10 mM Tris, pH 8.0, 150 mM NaCl, and 0.1% Tween 20). After blocking, the membranes were decorated with specific antibodies diluted in TBS-T as follows: PGR5 (1/2,500 dilution; Munekage et al., 2002), PGRL1 (1/10,000; DalCorso et al., 2008), NTA1 (1/500; (Li et al., 2023), PsaA (1/5,000; AS06 172), PsbE/F (1/10,000; AS06 112), PetA (1/1,000; AS20 4377), PetC (1/5,000; AS08 330), PC (1/10,000; AS06 141), PsaD1 (1/5,000; AS09 461) and cpFBPase1 (1/50,000; AS19 4319). Antibodies against PsaA, PsbE/F, PetA, PetC, PC, PsaD1 and cpFBPase1 were obtained from Agrisera (Vännäs, Sweden). Antibody against a peptide (CKSSSNNPEEPEKTQIQ) of PFSC1 (Q29Q44) was generated by BioGenes (Berlin, Germany) and used at 1/2,000 dilution in TBS-T. Proteins transferred onto the PVDF membrane were visualized by staining with Coomassie Brilliant Blue R-250 dye, ensuring equal loading of proteins. Signals were detected by enhanced chemiluminescence using the SuperSignal™ West Pico PLUS Chemiluminescent Substrate (ThermoFisher Scientific, Waltham, MA, USA) and a Fusion FX ECL reader system (Vilber Lourmat, Collégien, France). Signal intensities were quantified using ImageJ software (National Institutes of Health).

### Subcellular protein localization

The At2g04360 coding sequence without its stop codon was cloned into the pGWB5 vector (Nakagawa et al., 2007), and the vector was used to infiltrate extracted protoplasts from two-week-old *A. thaliana* seedlings as described previously (Dovzhenko et al., 2003).

### Alignment and phylogenetic tree

Multiple sequence alignment and phylogenetic tree were performed using the Geneious version 2023.0 created by Biomatters (https://www.geneious.com). Multiple sequence alignment was built using MUSCLE and homologous sequences to *A. thaliana* PFSC1 (Q29Q44 / NP_178517.2) identified using protein BLAST-NCBI (https://blast.ncbi.nlm.nih.gov/Blast.cgi), in different organisms: *Camelina sativa* (XP_019092803.1), *Brassica rapa* (XP_009142758.1), *Solanum lycopersicum* (XP_004243587.1), *Oryza sativa* (NP_001411979.1), *Lotus japonicus* (XP_057433373.1), *Anabaena sp. PCC 7120* (WP_010998185.1), *Synechococcus sp. PCC 7942* (WP_011243288.1), *Synechocystis sp. PCC 6803* (WP_010872027.1), *Leptolyngbya boryana* (WP_017286636.1), *Fischerella thermalis* (WP_102178976.1), *Chlorella variabilis* (XP_005851345.1), *Chlamydomonas reinhardtii* (XP_001700636.1), *Volvox carteri* (XP_002950523.1), *Monoraphidium neglectum* (XP_013906622.1), and *Ostreococcus tauri* (XP_003078581.2). The phylogenetic tree was constructed by maximum likelihood method, inferred using a neighbor-joining algorithm and Jukes-Cantor as the genetic distance model, using the Bootstrap (n= 100) method (Felsenstein, 1985).

### Proteomics analysis

Total proteins were extracted from 4-day-old WT and *pfsc1-1* seedlings (four biological replicates for each genotype, of which 1 WT and two *pfsc-1-1* were later excluded as outliers from further analysis). Proteome preparation, trypsin digestion, and liquid chromatography[tandem mass spectrometry (LC[MS/MS) were performed as previously described (Marino et al., 2019).

Raw data were processed using the MaxQuant software 1.6.17.0 (Cox and Mann, 2008), using the built[in Andromeda search engine (Cox et al., 2011) with default settings. Database searches were performed against the *A. thaliana* reference proteome (Uniprot, www.uniprot.org, version April 2021). Proteins were quantified across samples using the label[free quantification (LFQ) algorithm (Cox et al., 2014) with the match-between-runs option enabled. For downstream statistical analysis, Perseus version 1.6.15.0 (Tyanova et al., 2016) and RStudio version 1.2.5019 (Team, 2019) were used. Significantly differentially abundant protein groups were calculated employing the R/Bioconductor package limma (Andrews, 2010), with *P*-values adjusted for multiple comparisons according to the Benjamini−Hochberg approach (Benjamini and Hochberg, 1995). Proteins were considered to show a significant change with a fold change relative to the WT larger than 1.5 (up-regulated) or lower than 0.66 (down-regulated) and with an FDR < 0.05.

The mass spectrometry proteomics data were deposited at the ProteomeXchange Consortium via the PRIDE (Perez-Riverol et al., 2019) partner repository with the dataset identifier xxxxxxxx.

### Statistical analyses

Statistical significance of differences, except for the proteomics analysis, was assessed using one-way ANOVA with post-hoc Tukey HSD test (https://astatsa.com/, version October 2023).

### Accession numbers

ATG accession numbers: *AT2G05620* (*PGR5*), *AT4G19100* (*PAM68*), *AT4G03280* (*PGR1*) and *AT4G02770* (*PSAD1*). Additional accession numbers are listed in **Table 1**.

## Supplemental data

The following supplemental materials are available.

**Supplemental Figure S1.** Example of fluctuating light measurements.

**Supplemental Figure S2.** Alignments of the encoded DEIP1/NTA1 protein variants in WT, *p5s21*, *pgr5 deip1-Cas#1* and *pgr5 deip1-Cas#2* backgrounds.

**Supplemental Table S1.** Proteins expressed differentially in *pfsc1-1* compared to Col-0.

**Supplemental Table S2.** Oligonucleotide sequences used for genotyping, gRNA, sequencing and cloning.

**Supplemental Data Set 1.** List of all proteins identified by shotgun proteomics.

## ACKNOWLEDGMENTS

We thank Prof. Shikanai for providing *pgr5*, *pgr1* and *pgr5 pgr1* seeds and PGR5-specific antibody, and Prof. He for the NTA1-specific antibody. We also thank Hannah Kallus for her excellent technical assistance.

## AUTHOR CONTRIBUTIONS

Research was designed by D.L. and B.N.; J.F.P., B.N. and S.W. performed the experiments; G.M. performed the proteomics analysis; T.K. analyzed the resequencing data; D.L., B.N. and J.F.P. wrote the manuscript. All authors read and approved the final manuscript.

## SUPPLEMENTAL DATA

**Supplemental Table S1.**
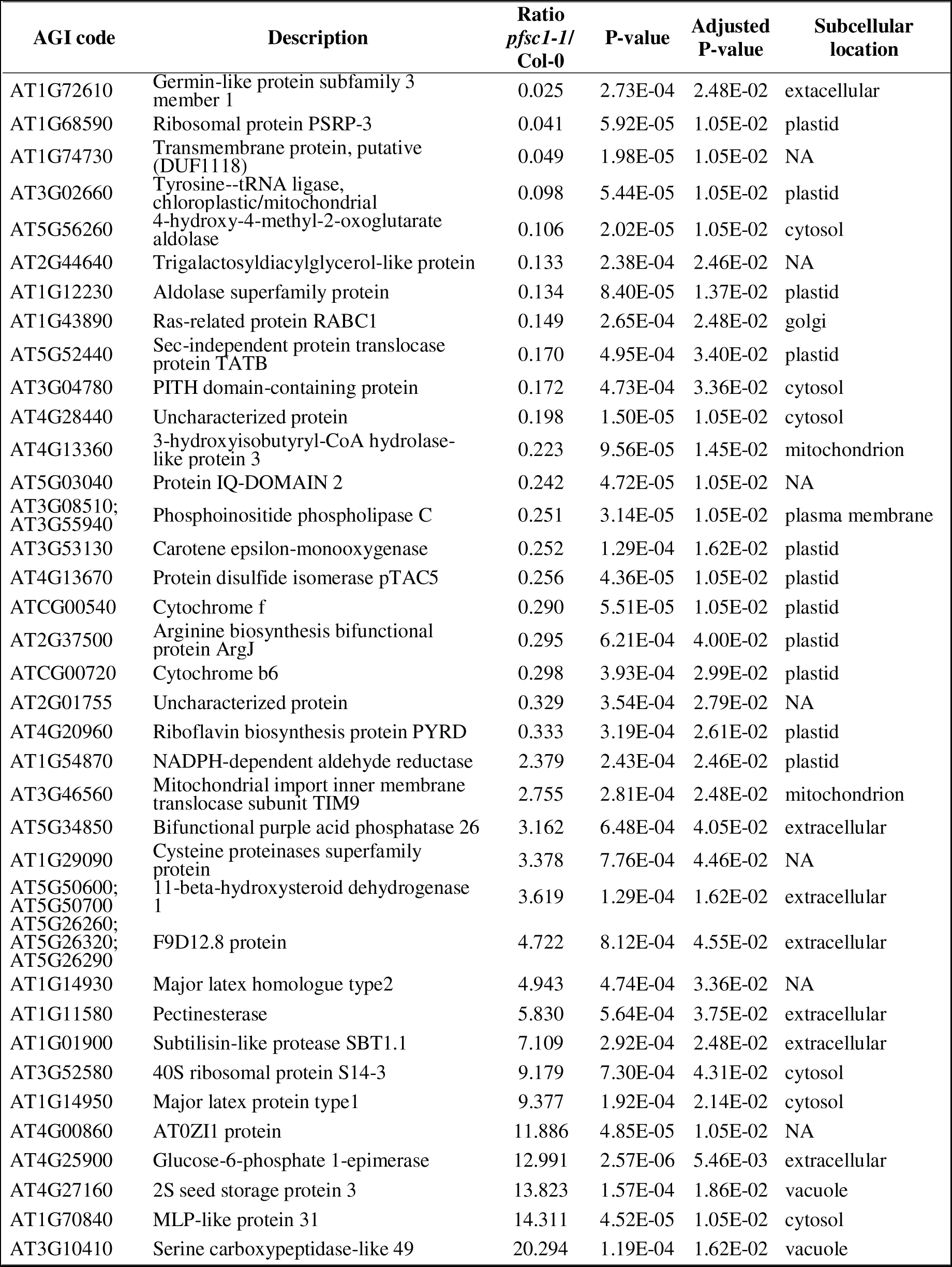
Proteins expressed differentially in *pfsc1-1* compared to Col-0.

**Supplemental Table S2.**
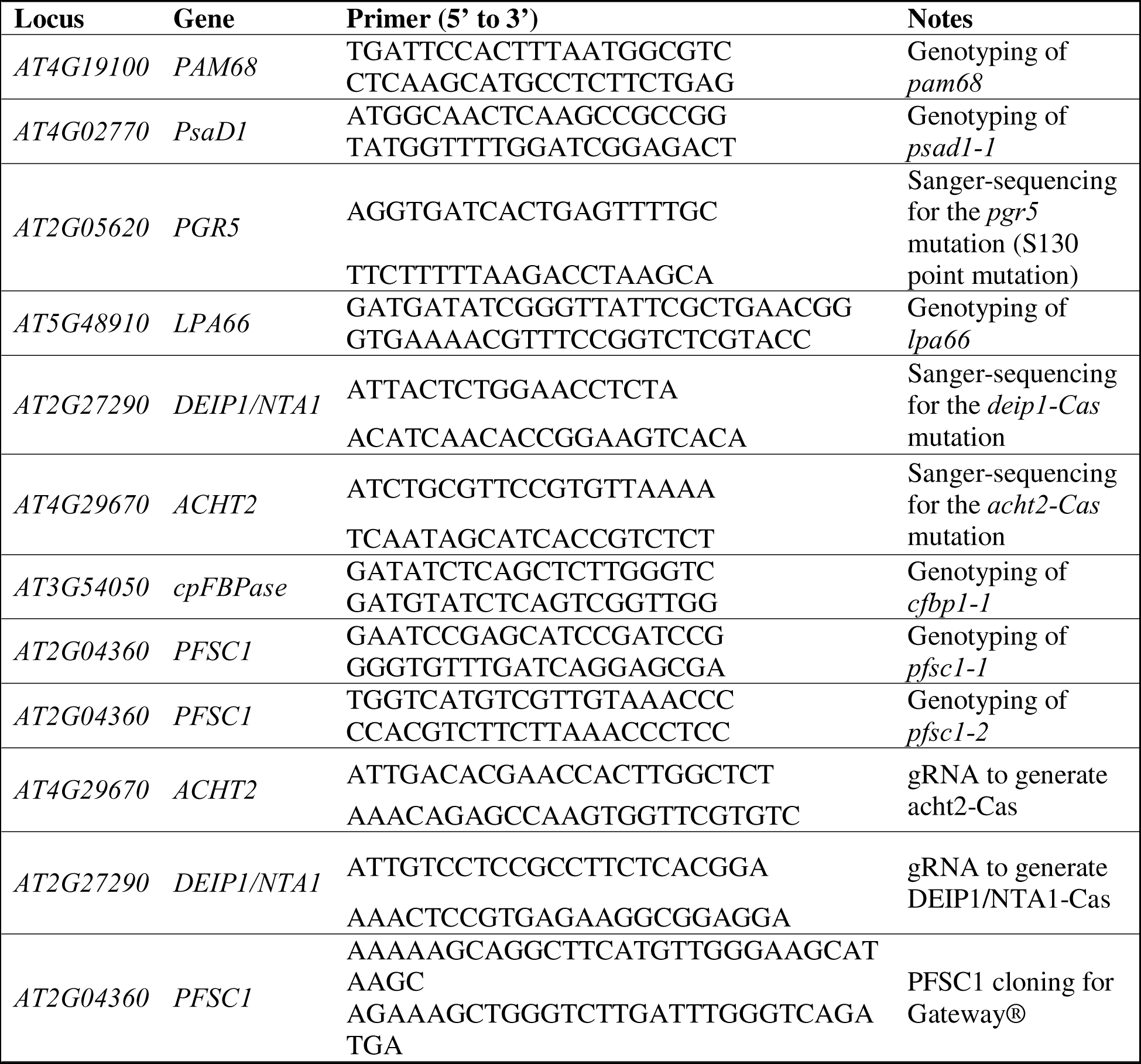
Oligonucleotide sequences used for genotyping, gRNA, sequencing and cloning.

**Supplemental Figure S1.**
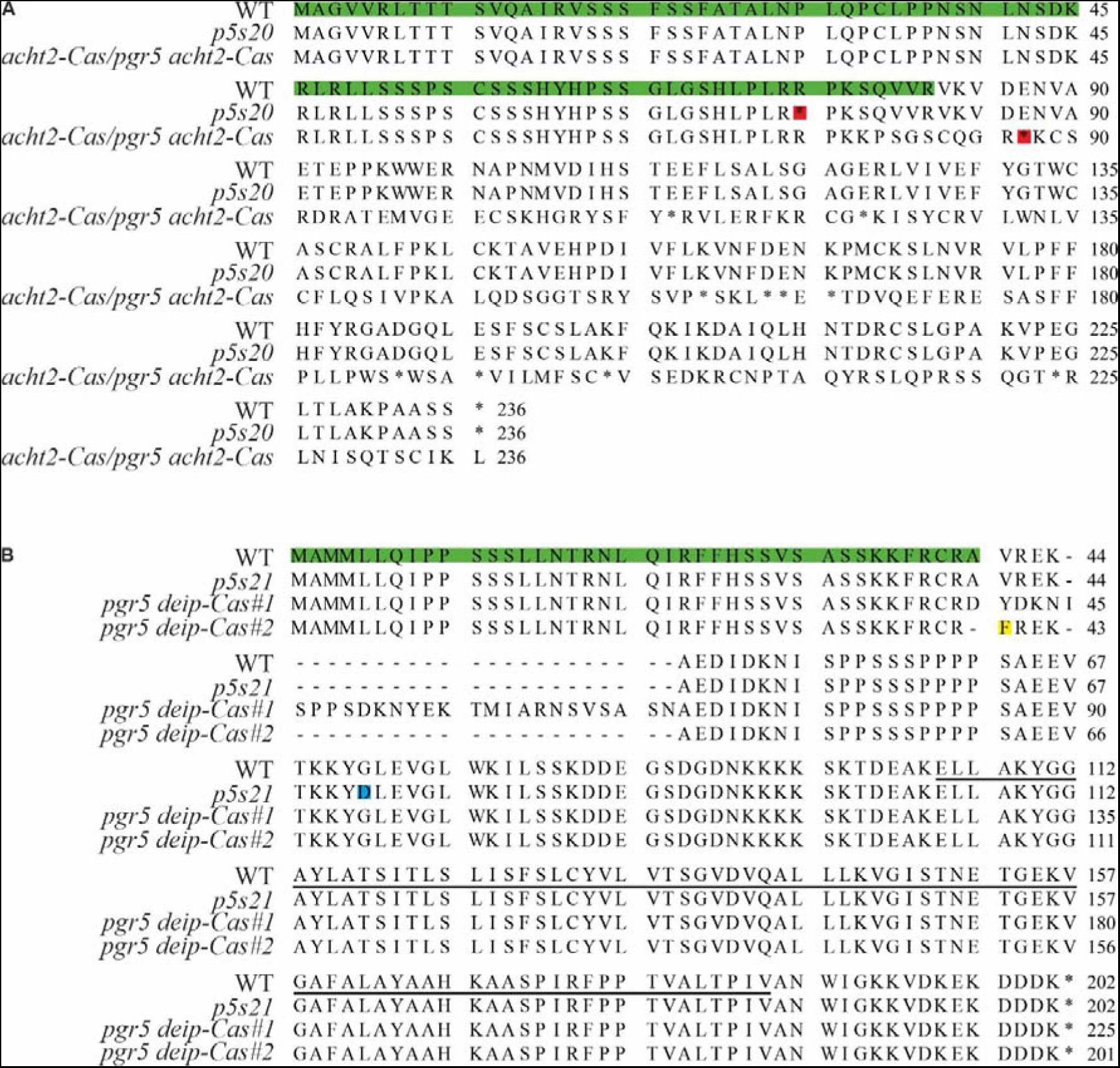
Mutations in proteins identified in the course of the *pgr5* suppressor screen and additional alleles generated by CRISPR-Cas. **(A)** Alignment of the ACHT2 protein variants in the WT (Col-0), *p5s20*, and *acht2-Cas/pgr5 acht2-Cas* backgrounds. The green background indicates the cTP, gaps are indicated by a dash. The red rectangle indicates the premature stop codon in the *p5s20* (at position 74) and *acht2-Cas /pgr5 acht2-Cas* (at position 86) line. The underlined amino acids correspond to the predicted transmembrane domain. **(B)** Alignment of the DEIP1/NTA1 protein variants in the WT (Col-0), *p5s21*, *pgr5 deip1-Cas#1* and *pgr5 deip1-Cas#2* backgrounds. Green background indicates the cTP, yellow background indicates the amino acid exchange in *pgr5 deip1-Cas#2.* Gaps are indicated by a dash. The blue rectangle indicates the G72D mutation in the *p5s21* line. The underlined amino acids correspond to the predicted transmembrane domain. Line *pgr5 deip1-Cas#1* showed an insertion of 69 nucleotides and line #2 a deletion of 3 as an additional nucleotide exchange, resulting in an insertion of 23 amino acids in the *pgr5 deip1-Cas#1* and to a deletion of one amino acid and an exchange from valine to phenylalanine at the position 40 in *pgr5 deip1-Cas#2*.

**Supplemental Figure S2.**
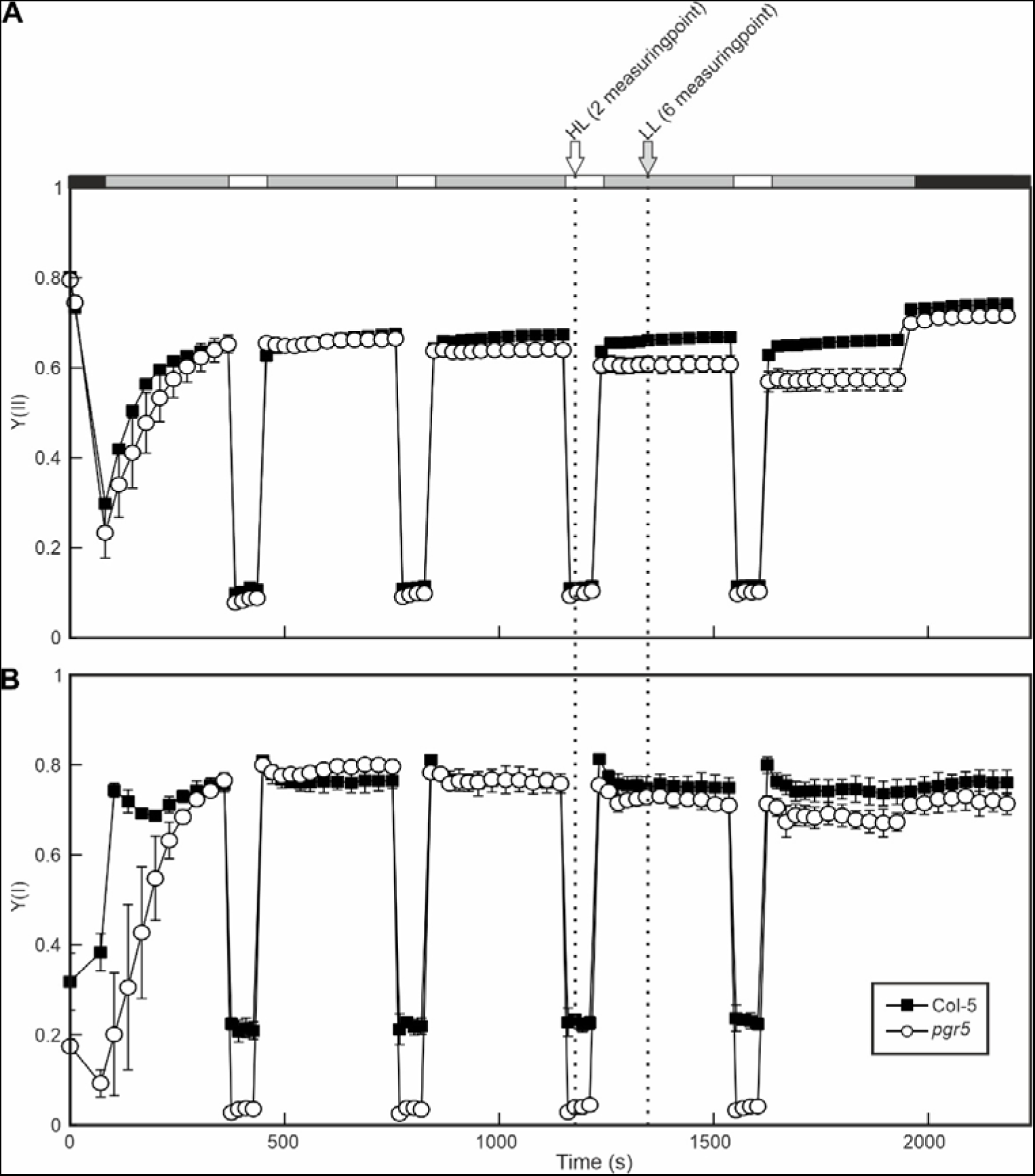
Example of FL measurements. Three-week-old WT (Col-5) and *pgr5* plants grown under control light conditions were subjected to a FL program using the DUAL-PAM chlorophyll fluorometer. This program mimics the conditions of the screen and consists of 5 cycles of 5 min of low actinic light (50 µmol photons m^−2^ s^−1^) alternating with cycles of 1 min moderately high actinic light (500 µmol photons m^−2^ s^−1^). The values corresponding to PSII (top panel) and PSI (bottom panel) quantum yields Y(II) and Y(I), respectively, show the average of at least 3 replicates ± SD.

